# DeepMol: An Automated Machine and Deep Learning Framework for Computational Chemistry

**DOI:** 10.1101/2024.05.27.595849

**Authors:** João Correia, João Capela, Miguel Rocha

## Abstract

The domain of computational chemistry has experienced a significant evolution due to the introduction of Machine Learning (ML) technologies. Despite its potential to revolutionize the field, researchers are often encumbered by obstacles, such as the complexity of selecting optimal algorithms, the automation of data pre-processing steps, the necessity for adaptive feature engineering, and the assurance of model performance consistency across different datasets. Addressing these issues head-on, *DeepMol* stands out as an Automated ML (AutoML) tool by automating critical steps of the ML pipeline. *DeepMol* rapidly and automatically identifies the most effective data representation, pre-processing methods and model configurations for a specific molecular property/activity prediction problem. On 22 benchmark datasets, *DeepMol* obtained competitive pipelines compared with those requiring time-consuming feature engineering, model design and selection processes. As one of the first AutoML tools specifically developed for the computational chemistry domain, *DeepMol* stands out with its open-source code, in-depth tutorials, detailed documentation, and examples of real-world applications, all available at https://github.com/BioSystemsUM/DeepMol and https://deepmol.readthedocs.io/en/latest/. By introducing AutoML as a ground-breaking feature in computational chemistry, DeepMol establishes itself as the pioneering state-of-the-art tool in the field.

## 1 Introduction

In recent years, computational chemistry has undergone a remarkable transformation, driven by advances in machine learning (ML) and deep learning (DL) techniques and by the immense growth of available chemical data [1, 2, 3, 4]. These have facilitated exploring and understanding the complex relationships between chemical structures and their properties [5]. As a result, using these computational methods within the chemical discovery pipeline has emerged as a powerful and promising way to expedite the discovery of new chemicals with improved properties [6].

Quantitative structure-activity/property relationship (QSAR/QSPR) models have always been a focal topic in computational chemistry [7, 8, 9, 10]. Initially, simple statistical models were applied to small datasets of molecules characterized by a restricted array of descriptors [11], offering a straightforward way to correlate molecular structure with biological activity [12]. Nowadays, QSAR/QSPR modelling consists of large sets of molecules with an extensive set of molecular descriptors.

As the amount of available data and the complexity of QSAR/QSPR models continue to grow, the demand for advanced techniques has led to the emergence of DL as a viable alternative to traditional ML models [13]. With the current plethora of ML/DL models and diverse chemical descriptors available, researchers face the challenge of selecting the most suitable alternative for their data regarding data representation and processing and ML/DL models [12]. Choosing the optimal combination of features/ descriptors and models requires exhaustive testing to comprehensively understand their performance on a given dataset [14, 15]. Recognizing this challenge, the importance of automated machine learning (AutoML) frameworks becomes unequivocal. In the fields of QSAR/QSPR modelling, where pre-processing steps, models, and their combination can significantly impact results, the need for a comprehensive and easily customizable AutoML framework is even more evident.

However, despite this need, only a few resources offer an easily customizable framework capable of providing a wide range of possibilities. ZairaChem [16] provides the first AutoML framework described in the literature that can automatically optimize all the pre-processing steps and model hyperparameters for specific QSAR problems. However, users cannot easily customize this system to suit their needs. For example, ZairaChem does not provide ways of creating custom objective functions, i.e. how to generate the final metric to give feedback to the AutoML system, e.g. whether in a cross- or hold-out validation or other scenarios. Another example is the impossibility of defining the type of models and descriptors to test. Furthermore, ZairaChem is restricted to binary classification tasks, which can be very limiting.

Herein, we present *DeepMol*, an AutoML python-based open-source framework for the prediction of activities and properties of chemical molecules. Although fully automated, *DeepMol* is built modularly, allowing for the customization of every step of the ML pipeline, starting from the data loading and processing to the model prediction and explainability. Nonetheless, what sets DeepMol apart is its robust AutoML functionality, allowing the automatic optimization of different scenarios involving preprocessing methods, data engineering techniques, and ML/DL models and respective hyperparameters. Even for users with minimal coding experience, with just a few lines of code, *DeepMol* allows testing thousands of configurations to determine the most effective ones for their specific datasets. Notably, *DeepMol* provides documented ways of defining objective functions and customizing AutoML experiments. Moreover, it is suitable for several ML tasks such as binary, multi-class, multi-label/multi-task classification and regression. The framework leverages other well-known and established packages, like *RDKit* [17] for molecular operations, *Scikit-Learn* [18], *Tensorflow* [19] and *DeepChem* [20] for model building and *Optuna* [21] for end-to-end ML pipeline optimization. *DeepMol* is available at https://github.com/BioSystemsUM/DeepMol. Additionally, it is easily installed through PyPi, conventionally installed with *pip install deepmol[all]*, or as a docker image *docker pull biosystemsum/deepmol:latest*.

Comprehensive documentation with examples for each step described in this paper is provided at https://deepmol.readthedocs.io/en/latest/.

A rigorous experimental framework was established to ensure an impartial assessment of the AutoML engine’s capabilities, enabling evaluation of its performance across 22 benchmark datasets for predicting adsorption, distribution, metabolism, elimination, and toxicity (ADMET) derived from the Therapeutics Data Commons (TDC) repository [22]. All the models can be accessed in https://doi.org/10.5281/zenodo.11184008 and the code for the experiments in https://github.com/ BioSystemsUM/deepmol case studies.

## 2 Implementation

*DeepMol* was primarily developed as a comprehensive end-to-end AutoML framework for computational chemistry. Yet, its modular design permits the independent utilization of its components. The framework provides a wide range of techniques for each step of a general ML pipeline, from data pre-processing and feature extraction to model training, evaluation, and explainability. Furthermore, the package has been extended with additional functionalities of significant relevance, including unsupervised learning models, data-splitting strategies specifically designed for molecular datasets, and approaches to address data imbalance.

The *DeepMol* AutoML engine, as illustrated in Figure 1, comprehensively explores a vast array of potential methodological combinations. In this context, state-of-the- art computational chemistry methods are employed automatically and sequentially in a pipeline, following a specific configuration, ensuring it is tailored to address specific chemical tasks. The configuration space encompasses several data standardization methods (Figure 1a: three methods), feature extraction methods (Figure 1b: four options for sets with 34 methods in total, all with their respective parameters), scaling and selection methods (Figure 1c: 14 methods and respective parameters), various ML models and ensembles (Figure 1d: 140 models), and their respective hyperparameters. Upon starting the AutoML experiment, the engine first processes the training data following a predefined sequence of steps, known as the pipeline configuration, and then uses this data to train an ML/DL model. Post-training, a separate set of data, the validation set, is processed and assessed to evaluate the model’s performance. The outcomes of this evaluation are then fed back into the optimization framework, guiding it in choosing new parameters and methods for improvement. This cycle of training and evaluating is repeated a user-specified number of times, referred to as *trials*. After completing all the trials, the system analyzes the results to identify and select the most effective pipeline. The optimization framework is efficiently powered by the *Optuna* search engine, which provides nine different optimization algorithms compatible with *DeepMol*. Once the optimal pipeline has been identified, it can be applied to analyze new data, enabling predictions on untested data. Additionally, this pipeline facilitates the virtual screening of extensive databases, efficiently identifying potential molecules of interest.

**Fig. 1.**
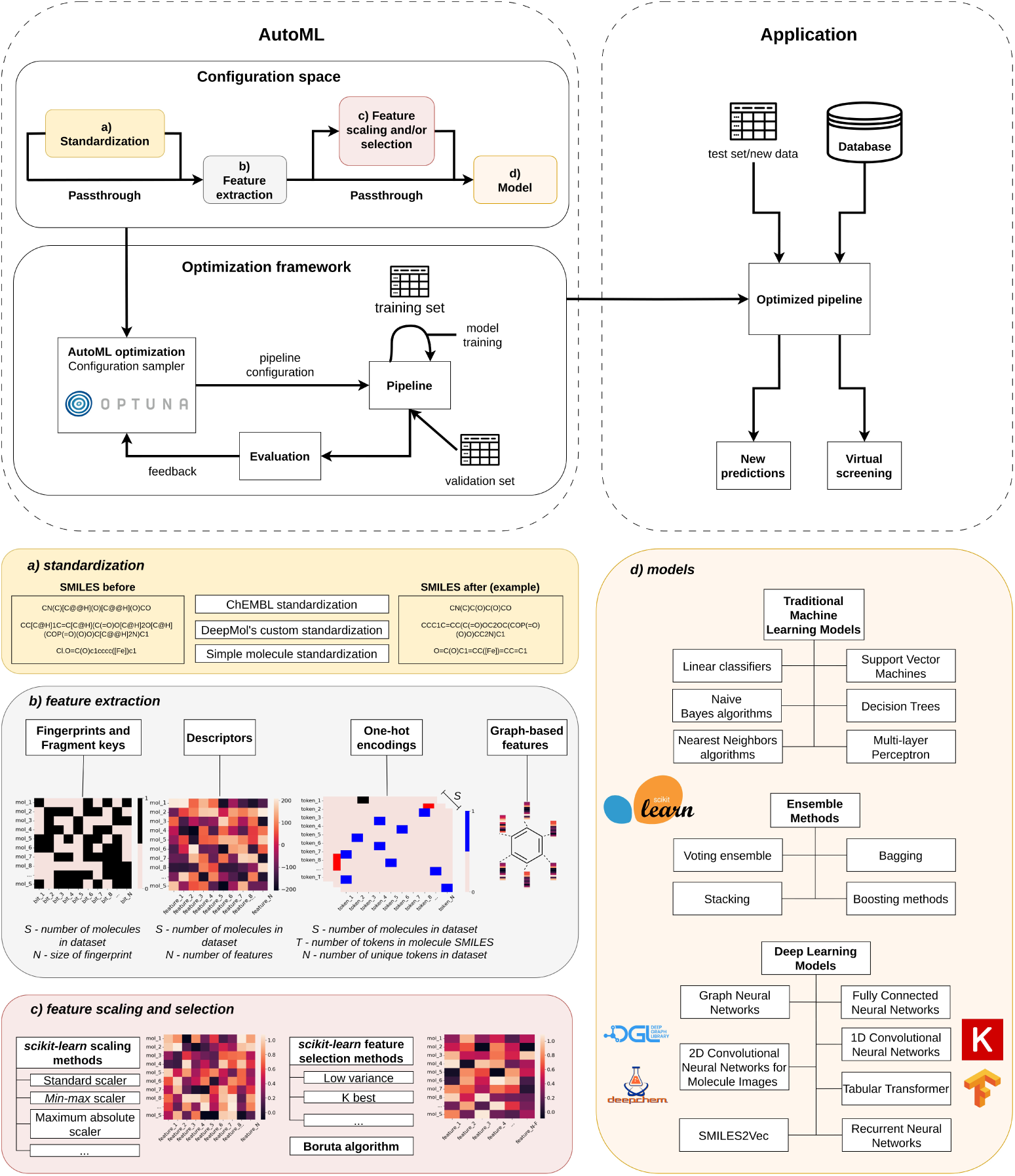
The DeepMol AutoML framework offers a comprehensive approach that includes an optimization framework that samples pipeline configurations from the configuration space. This pipeline begins with an optional **(a)** standardization step (configuration space: 3 different methods), followed by **(b)** the extraction of features (configuration space: four options for sets with 34 methods in total, all with their respective parameters). These features can then be optionally **(c)** scaled and selected (configuration space: 14 methods and respective parameters), preparing them for **(d)** the training phase of the model (configuration space: 140 models, architectures and hyperparameters). Post-training, the model’s performance is evaluated based on a predetermined metric. This evaluation feedbacks to the *Optuna* optimization engine. *Optuna* uses these results to inform its selection of new parameters and methods for enhancing the pipeline’s efficiency using state-of-the-art optimization algorithms. This process is repeated *n* times (chosen by the user), also called trials, and, in the end, the best pipeline is selected. Each individual box represented in the configuration space is expanded and schematized in the boxes below. Finally, the result of the AutoML engine is an optimized pipeline that can be used to transform new data and perform new predictions.

This section provides an overview of the key components and capabilities of *Deep-Mol*, highlighting its significant contributions and advantages in chemical compound analysis.

### 2.1 Data Loading

The representation of molecular structures can be achieved using various formats. One such format is the Simplified Molecular Input Line Entry System (SMILES), which is readable and concise, as short as single-line ASCII strings [23]. The other format is a connection table, which can be stored as a Structure Data File (SDF) and used to represent three-dimensional structures of molecules [24, 25]. These are the *de facto* standard starting points for representing molecules in ML tasks [23, 26] and *DeepMol* provides loaders for both formats. Once loaded, the information is converted into a structured data format that allows easy access to relevant details, such as the SMILES representation of all molecules, their identifiers, and known outputs associated with the input data (labels). Alternatively, if one prefers to load molecules from an already-loaded data frame, a *SmilesDataset* class can be created. This dataset accepts SMILES or *RDKit Mol* objects as input, along with their corresponding identifiers and labels.

### 2.2 Molecular Standardization

The availability of vast amounts of chemical data found in molecular databases containing hundreds of millions of compounds can be a double-edged sword. While it pushes the field forward, it makes the human curation process infeasible, resulting in the frequent occurrence of incorrect and inconsistent molecular structures [27]. Even minor structural errors and inconsistencies within a dataset can result in significant losses in the predictive ability of QSAR/QSPR models [28]. Consequently, the development of strategies that can tackle this problem has received increased attention in recent years [29, 30, 31].

With *DeepMol*, it is possible to standardize molecules using three different options:

- *BasicStandardizer* performs basic sanitization using *RDKit* to ensure that a molecular structure is represented consistently and validly according to a set of predefined rules. These include kekulization, check of valencies, set aromaticity, conjugation, and hybridization.
- *CustomStandardizer* performs a set of steps defined by the user, using *RDKit*. Some steps include molecular sanitization (same as in the BasicStandardizer), removal of isotope and/or stereochemistry information, neutralizing charges, fragment removal, and kekulization.
- *ChEMBLStandardizer* performs a set of rules as used by the ChEMBL database [29]. It consists of three components: a Checker that tests the validity of the chemical structures, a Standardizer that formats compounds based on the U.S. Food and Drug Administration (FDA) and International Union of Pure and Applied Chemistry (IUPAC) guidelines; and a *GetParent* component that removes any salts and solvents from the compound.

### 2.3 Feature extraction

In ML, feature extraction from molecules is a common task. Different types of molecular features are categorized from 0-dimensional (D) to 4D. 0D features provide overall information about the molecule (e.g. atom and bond counts), while 1D features describe substructures within the molecule (e.g. fingerprints) [32, 33]. 2D features capture molecular topology, and 3D features capture geometric information of the molecule’s three-dimensional structure [34]. 4D features are descriptors with an additional dimension representing interactions with receptors’ active sites or multiple conformational states.

*DeepMol* provides a set of 0D, a few 1D and 2D descriptors in only one class. The descriptors contained in this class are enumerated and described in Supplementary Table S1. *DeepMol* offers a wide range of 1D features provided by *RDKit*, including circular, atom pair, layered and *RDKit* fingerprints. *DeepMol* also includes Molecular ACCess System (MACCS) keys implemented by *RDKit*, which encode the presence or absence of specific molecular fragments or substructures in a molecule. These are further enumerated and described in Supplementary Table S2.

3D molecular conformations must be generated or loaded before generating 3D descriptors. For this matter, *DeepMol* provides conformer generation methods like Experimental-Torsion basic Knowledge Distance Geometry (ETKDG) and methods like Merck Molecular Force Field (MMFF) and Universal Force Field (UFF) for optimizing the generated conformers. Once generated, *DeepMol* offers methods to extract features from these conformers. These methods include AutoCorr3D, Radial Distribution Function (RDF), the plane of best fit, MORSE, WHIM descriptors, Radius of Gyration, shape descriptors, and principal moments of inertia. More details on these methods can be found in Supplementary Table S3.

*DeepMol* also provides one-hot encoding schemes. Herein, *DeepMol* provides an implementation of an atom-level tokenizer and *k* -mer tokenizer from Li et al. [35]. The former method permits treating each character in the SMILES string linked to an atom (e.g., [C@] or [N@+]) as an identical token, whereas the latter provides groups of *k* characters of the SMILES string. Both create a vocabulary dynamically given a dataset. If no tokenizer is passed as input, the one-hot encoding uses the atom-level tokenizer.

*DeepMol* extends *DeepChem* to provide inputs for graph neural networks (GNN), including molecular graphs (2D descriptors), representing the molecule by a list of neighbors and a set of initial feature vectors for each atom. The feature vectors represent the atom’s local chemical environment, including atom types, charge, and hybridization, among others [36]. Other implementations for using GNNs were also included, such as methods for implementing Duvenaud graph convolutions [37], Weave convolutions [36], and MolGAN [38]. Coulomb matrices and their eigenvalues [39] are also provided as methods of extracting 3D features, providing information on electrostatic interaction between atoms.

Finally, *DeepMol* also extends *DeepChem* to convert molecules into images and encodings to be passed to an embedding layer. All DeepChem-derived feature extractors are further detailed in Supplementary Table S4.

### 2.4 Data Scaling

Many feature extraction methods encode molecules as bit vectors, while others use vectors of real numbers. In the latter, a challenge arises when the features have different numerical ranges. This discrepancy can cause certain features to dominate the learning process due to their larger scales. Data scaling becomes crucial to address this issue. Bringing all features into a similar numerical range can enhance the performance and convergence of many ML algorithms. *DeepMol* offers a wide range of data scalers, as provided in the *Scikit-Learn* package (Supplementary Table S5).

### 2.5 Feature Selection

Feature selection is a common task in ML problems. By choosing the most relevant features, the performance of an ML model can be significantly enhanced. Eliminating irrelevant or redundant features helps prevent overfitting and improve the generalization power of the model, while reducing the computational burden. *DeepMol* supports many types of feature selection as provided by *Scikit-Learn*, including removing features with low variability, supervised filters based on statistical tests, as well as wrappers such as Recursive Feature Elimination, or selecting features based on importance weights (Supplementary Table S6). In addition, *DeepMol* also provides feature selection based on the Boruta algorithm [40].

### 2.6 Machine and Deep Learning models

*DeepMol* offers compatibility with three popular ML/DL frameworks: *Tensorflow*, *Scikit-Learn*, and *DeepChem*. This allows seamless integration of any model built upon these frameworks into *DeepMol*. A wide range of pre-built models from these frameworks is readily available, supporting single and multi-task problems, binary and multi-class classification and regression, making it convenient for users to utilize these models for specific tasks.

Through *Scikit-Learn*, *DeepMol* offers an extensive selection of popular ML models, including, among others, linear and logistic regression, support vector machines, decision trees, random forests, and gradient boosting. In conjunction with *DeepChem*, it also provides several DL models specifically tailored for chemical data, including graph neural networks (GNNs), recurrent neural networks (RNNs), and transformer models. Moreover, the flexibility offered by *Tensorflow* enables integration of any DL architecture into *DeepMol* ’s pipeline. By consolidating these capabilities, *DeepMol* serves as a comprehensive and versatile framework, facilitating implementation and comparison of various ML and DL models in one unified platform [41, 42].

### 2.7 Hyperparameter Optimization

In *DeepMol*, users can perform hyperparameter optimization for all the abovementioned models. This can be done either by using a validation dataset or through cross-validation. Fine-tuning the model’s hyperparameters is crucial to obtaining optimal results, mitigating overfitting, and improving overall efficiency. Currently, *DeepMol* utilizes *randomized* or *grid search* methods to explore and find the best hyperparameters.

### 2.8 Pipelines

In *DeepMol*, we have introduced a pipeline class that simplifies the creation of an ML pipeline. This class empowers users to construct an ML pipeline tailored to their specific needs. It offers the flexibility to incorporate a diverse range of steps, enabling the inclusion of any combination of the following methods:

- Data standardization;
- Feature extraction or transformation (data scaling and selection);
- Model training and hyperparameter tuning.

Pipelines in *DeepMol* work the same as *Scikit-Learn* pipelines: use the *fit transform* method to fit and transform the train data and the *transform* method for solely transforming the validation and test sets. The pipeline class also allows users to save and load the pipeline again. Moreover, it allows to evaluate the pipeline and make predictions on new data using the *evaluate* and *predict* methods, respectively. The model herein declared can also be optimized.

Users can design and customize their ML pipeline, adapting it to their unique requirements and desired outcomes. This approach significantly reduces the complexity of constructing an ML pipeline, promoting efficient experimentation and development.

### 2.9 AutoML: pipeline optimization

Pipelines provide a convenient and efficient way to transform raw data into trained and validated models, but determining the appropriate sequence of steps often requires expertise. Moreover, if one wishes to explore multiple combinations of pre-processing techniques (data *Transformers*) and models (*Predictors*), manually constructing a *Pipeline* for each combination can be repetitive and time-consuming.

Automating the process of building and optimizing each step and respective parameters of a *Pipeline* can greatly assist researchers with limited ML expertise in building and deploying effective QSAR/QSPR systems, accelerating the development process. In *DeepMol*, we offer the *PipelineOptimization* module (AutoML), which enables the search for the best configurations from a large number of possibilities. This module leverages the powerful *Optuna* library [21] and its state-of-the-art optimization algorithms.

As with any optimization problem, pipeline optimization needs an objective function. By default, *DeepMol* provides the objective of maximizing or minimizing a metric (e.g. accuracy) for a given validation set. However, other custom objective functions can be added.

Furthermore, a function can be created with the configuration space that will be optimized by *Optuna*. This function will receive a *Trial* object from *Optuna* and will return a set of steps (i.e. *Transformers* and *Predictors*) from the configuration space to define the trial pipeline. Alternatively, *DeepMol* provides a set of predefined configuration spaces (pre-sets) for convenience. The available pre-sets include:

- ’sklearn’: optimizes between all available Standardizers, Featurizers, Scalers, Feature Selectors in *DeepMol*, and all available models in *Scikit-Learn*.
- ’deepchem’: optimizes between all available Standardizers in *DeepMol* and all available models in *DeepChem* (depending on the model, different Featurizers, Scalers and Feature Selectors may also be optimized).
- ’keras’: optimizes between Standardizers, Featurizers, Scalers, Feature Selectors in *DeepMol*, and some core Keras models (Fully Connected Neural Networks (FCNNs), 1D Convolutional Neural Networks (CNNs), RNNs, Bidirectional RNNs, and transformer models).
- ’all’: optimizes between all the above.

Finally, a *Voting Pipeline* composed of the best five pipelines is created after running the AutoML, where the voting process for classification occurs through soft and hard voting mechanisms. In soft voting, the predicted probabilities from each classifier are averaged to determine the final prediction. On the other hand, hard voting involves a simple majority vote, where the class predicted by the majority of the individual classifiers is selected. Additionally, for regression tasks, the final prediction is obtained by averaging the predictions from each regressor, providing a robust ensemble prediction from the diverse models in the voting pipeline.

### 2.10 Unsupervised Learning

Unsupervised learning has become increasingly important in exploring the vast amounts of available and sometimes unlabeled chemical data. Through techniques like clustering and dimensionality reduction, unsupervised learning allows the identification of patterns and relationships in large datasets without prior knowledge.

*DeepMol* provides a simple interface to various techniques, such as Principal Component Analysis (PCA), t-distributed Stochastic Neighbor Embedding (t-SNE), K-means clustering, and Uniform Manifold Approximation and Projection (UMAP). These techniques can help scientists to efficiently analyze and visualize complex molecular data, providing valuable insights to guide subsequent analyses.

### 2.11 Data splitting

Splitting the data into training, validation and test data sets or in a cross-validation setting is a crucial step in ML pipelines. It is common for practitioners to require data separation based on molecular structures or similarities. Practitioners may opt for a homogeneous data split to maintain a balanced distribution of molecular structures or similarities across the sets. This approach aims to capture the diversity of the dataset, while providing reliable performance estimates for the ML model. Alternatively, practitioners may emphasize the separation of highly similar molecules from less similar ones (heterogeneous splits). By isolating similar molecules, practitioners can better assess the model’s ability to capture nuanced differences and generalize to unseen data. *DeepMol* provides molecular splitters that split the inputted dataset based on similarity, scaffolds or Butina clusters [43], while maintaining the stratification of classes. Figure 2 illustrates how the similarity and scaffold splits are distributed in the overall chemical space and how the homogeneity parameters control the split.

**Fig. 2.**
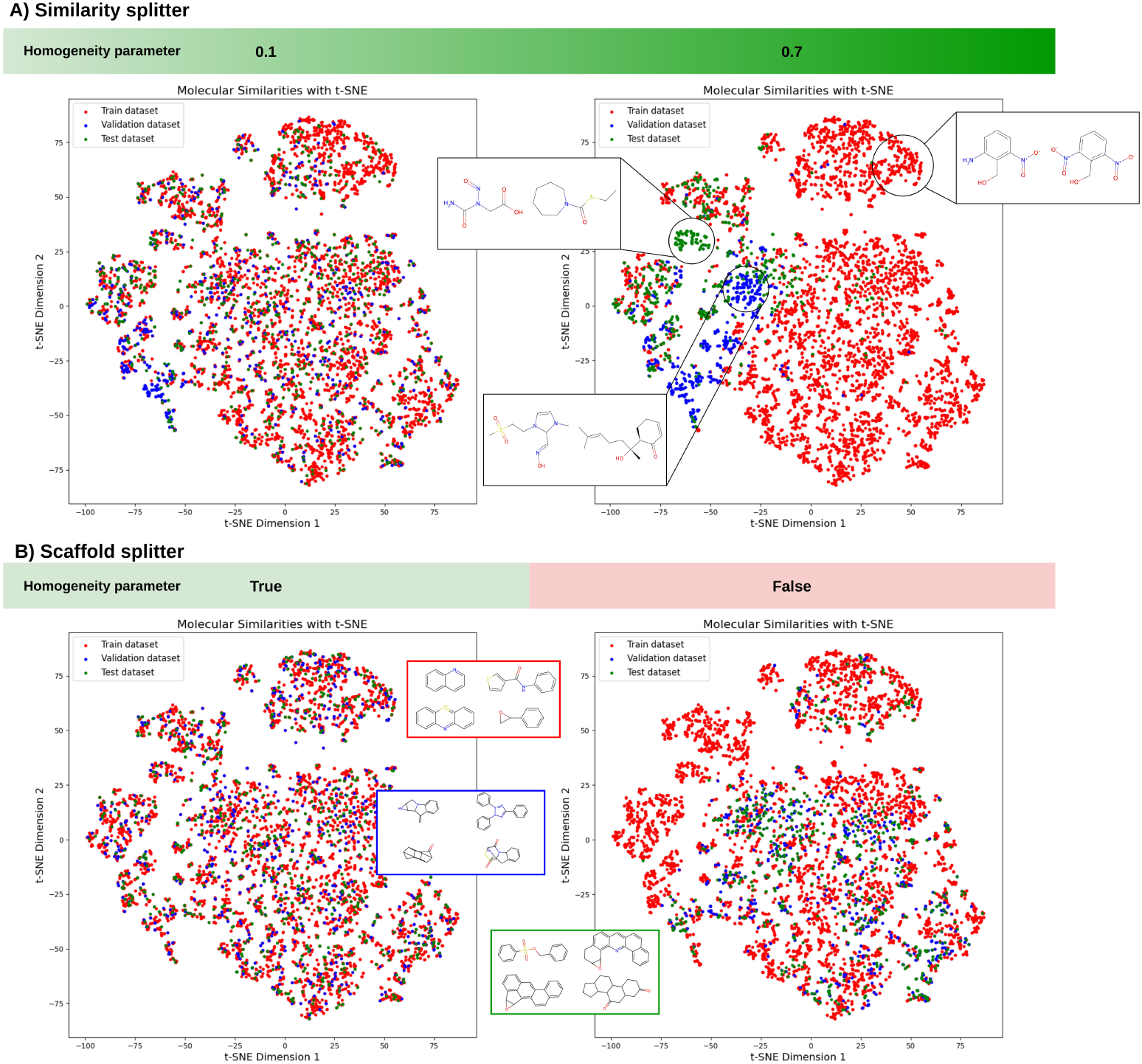
Depiction of dataset splits in the chemical space for the mutagenicity dataset from [44] provided by TDC. A similarity matrix was generated using the Tanimoto similarity metric over Morgan fingerprints. A t-SNE was applied to the matrix to reduce dimensionality and visualization. **A)** represents the similarity splitter and examples of molecules belonging to small clusters. One can regulate the homogeneity of the splits by tweaking the homogeneity parameter, which considers all the compounds with a similarity lower than the homogeneous parameter to be in the same set. The higher the parameter is, the more heterogeneous the split will be. **B)** represents the scaffold splitter and scaffolds belonging to the splits on the plot on the right. This splitter can separate the data by putting compounds with the same scaffolds in different data splits (homogeneous split) or not (heterogeneous split).

Moreover, *DeepMol* provides random and stratified splitters for single and multitask classification problems. More information about each splitter is provided in Supplementary Table S7.

### 2.12 Imbalanced data

Imbalanced data is a common problem in many real-world applications that heavily affect the quality and reliability of ML and DL approaches [45]. Likewise, imbalanced data is an issue in QSAR studies, where the difference between the number of active and inactive molecules can be extreme [46]. This difference generally leads to biased and sub-optimal models, as traditional ML algorithms may not be able to learn the minority class effectively.

Data balancing techniques have shown potential in mitigating the effect of imbalanced data on the model performance [46]. *DeepMol* provides various methods for handling imbalanced data, including over-sampling, under-sampling, and combination methods, as in the *imbalanced-learn* package [47]. For over-sampling, it provides a random over-sampler and Synthetic Minority Over-sampling Technique (SMOTE) [48], while Random under-sampler and *ClusterCentroids* can be used for under-sampling. Finally, for simultaneous over and under-sampling, it provides SMOTE with Edited Nearest Neighbours (SMOTEENN) [49] and SMOTE using Tomek links (SMOTETomek) [50]. More information about each imbalanced learning technique is provided in Supplementary Table S8.

### 2.13 Feature importance and model interpretability

Model interpretability, an essential aspect in the field of ML, refers to the capacity to comprehend and elucidate the internal mechanisms of an ML model. This ability provides insights into the rationale behind the model’s predictions or decisions, fostering a sense of trust in the model’s outputs.

SHAP (SHapley Additive exPlanations) [51] stands out as a widely utilized technique for achieving model explainability. It leverages the Shapley value, a fundamental concept derived from cooperative game theory, to assign significance to each input feature within a model. By comparing the predictions of various feature subsets, both with and without a specific feature, the Shapley value quantifies the marginal contribution of that feature to a prediction.

Visualizing SHAP values can take various forms, including bar charts, scatter plots, and summary plots. These visual representations facilitate the interpretation of how individual features influence the model’s predictions, aiding in identifying the most crucial features for a given prediction scenario.

Along with these methods, *DeepMol* allows one to visualize the structures associated with bits in Morgan and *RDKit* fingerprints and MACCS keys. Therefore, after computing the SHAP values, one can directly cross information from the most relevant features to the molecule and draw biological and chemical conclusions, as depicted in Figure 3.

**Fig. 3.**
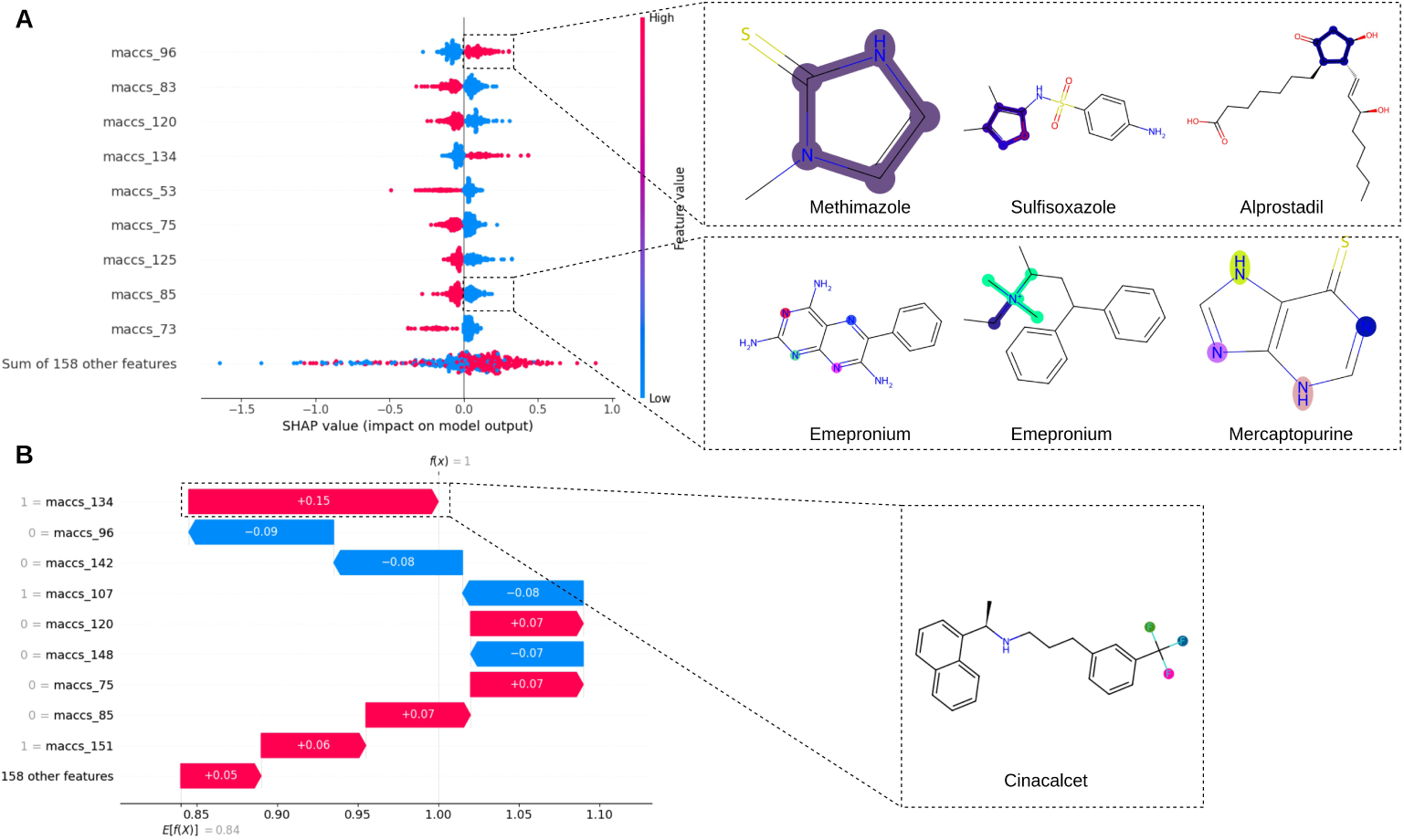
Depiction SHAP values for a ridge classifier and the most relevant MACCS keys in the molecules. The dataset used was the bioavailability dataset from [52] provided by TDC. **A)** provides the overall feature importance of all labels for all data points and depicts the two of the most relevant MACCS keys for the prediction. The highlighted atoms and bonds are the ones that correspond to the MACCS key. **B)** provides a depiction of the most important features of the drug cinacalcet, showcasing how it is also possible to assess features’ importance for individual cases.

## 3 Results

### 3.1 Results on the ADMET benchmarks

In this section, we discuss the performance of DeepMol AutoML on the 22 TDC ADMET benchmark group datasets. Thousands of pre-processing and feature extraction steps, models, and respective hyperparameters were tested and optimized using a fully automated process.

Each benchmark dataset within TDC includes its own training, validation, and test sets, alongside an evaluation metric. Out of the 22 datasets used, a combination of binary classification and regression tasks is covered. The dataset splits are computed through a scaffold split method, ensuring that the training and test sets feature distinct molecular structures. For regression tasks, the primary evaluation metric was the mean absolute error (MAE), with some exceptions where Spearman’s correlation (SC) is employed. Binary classification datasets were evaluated using the area under the receiver operating characteristic (AUROC) metric when the balance between positive and negative examples was even. Alternatively, when negative examples substantially outnumber positive ones, the area under the precision-recall curve (AUPRC) was used.

Benchmarking involves assessing an ML model’s performance using specific ADMET datasets. This is done by first dividing these datasets into training, validation, and test sets using five distinct data seeds to ensure variability. The model is then trained using the training and validation sets. Its effectiveness is evaluated based on how it performs on the test set. Finally, the process wraps up by calculating the average and standard deviation of the model’s predictions across all five seeds, providing a comprehensive measure of its reliability and accuracy.

Our experimental setup involved using the DeepMol AutoML engine to train models with the training sets and evaluate them on the validation sets. The optimization criterion for the *Optuna* search engine was to optimize the mean of the selected metric values across all validation sets. The Tree-structured Parzen Estimator (TPE) algorithm was utilized across 100 trials for hyperparameter and method selection.

Upon the conclusion of each experiment, the optimal pipeline was identified following the objective function criteria. Subsequently, the model underwent retraining by integrating both the validation and training datasets and its performance was assessed on the test set in alignment with the guidelines specified for submission. *Voting Pipelines* are also included in this analysis.

We compared our results with the TDC ADMET benchmark group leaderboard submissions. Overall, our AutoML framework demonstrated good performance, securing top-1 placement in one benchmark and achieving top-3 and top-5 rankings in four and eleven benchmarks, respectively. While our performance may not have been outstanding across all benchmarks, it is crucial to understand the remarkable nature of these results. This is particularly noteworthy, considering that the majority of benchmark submissions typically rely on handcrafted pre-processing steps and models selected for their performance. Our AutoML framework, operating without explicit instructions and requiring no prior domain knowledge from the users, efficiently navigated an extensive array of possibilities. That resulted in fully reproducible end-to-end pipelines that demonstrated competitive results. More details on each of the voting pipelines and ensembles are given in Supplementary Table S11.

#### 3.1.1 Absorption benchmarks

The *absorption benchmarks* measure how a drug travels from the administration site to its action site. These benchmarks consist of six datasets, three of which involve classification, and three involve regression. The AutoML framework of DeepMol achieved the top rank on the leaderboard for the human intestinal absorption (HIA) dataset and the second position for the aqueous solubility (AqSol) dataset. Interestingly, the HIA dataset, comprising 578 molecules, is the smallest in the benchmark, while the AqSol dataset, consisting of 9982 molecules, is the largest. This shows the effectiveness of our framework in finding robust models across varying data sizes. Additionally, it is noteworthy that graph-based models outperformed others in three out of the six benchmarks. The results for the six benchmarks and respective pre-processing steps and models can be seen in Table 1.

**Table 1.**
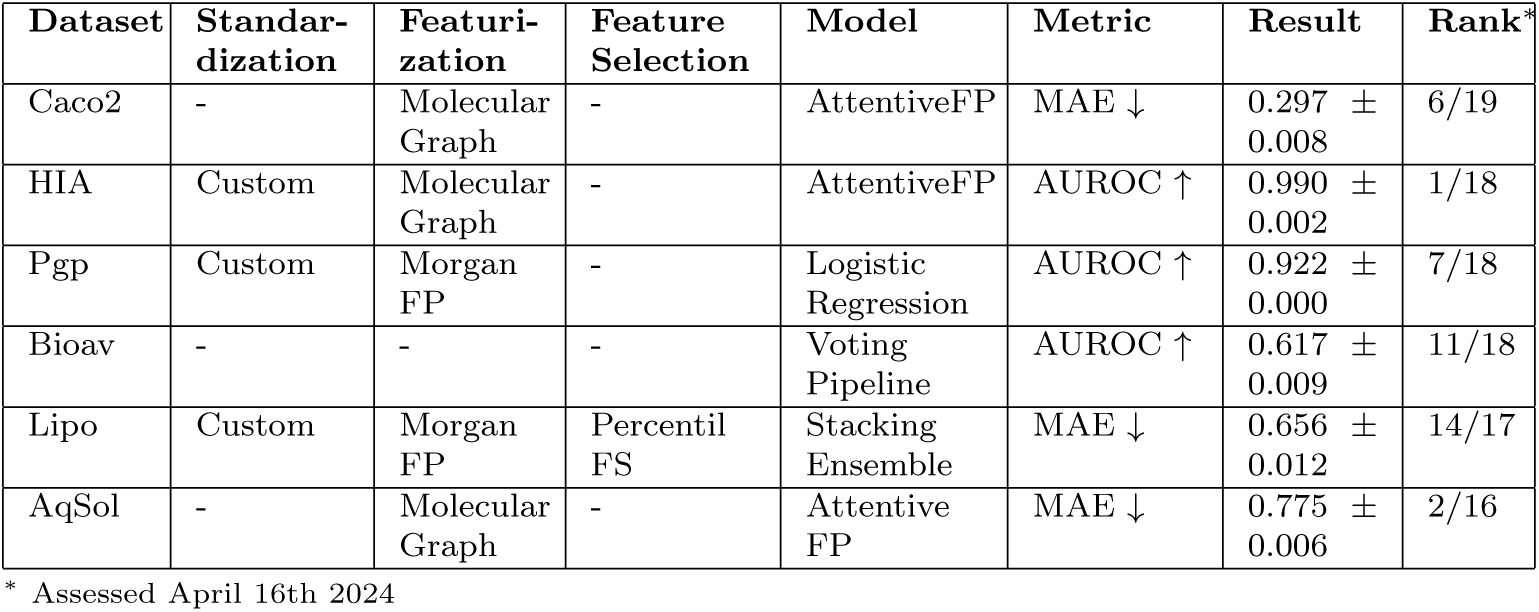
Performance of DeepMol in the six absorption-related properties in the TDC ADMET benchmark group.

Although DeepMol did not provide so much competitive performance for the drug oral bioavailability (Bioav) and lipophilicity (Lipo) datasets, the AutoML results provided invaluable hints regarding the best-performing methods. *Optuna* provides a dashboard to explore the results across all the trials. We analysed the results and focused our pipeline optimization on the best-performing methods and ML models. This allowed us to identify Histogram-based Gradient-boosting classification trees as top-performing models for the Bioav dataset and stacking ensembles for the Lipo dataset, leading us to set up smaller experiments and reduce the configuration space of the method search. We improved the AUROC on the Bioav test set from 0.617 to 0.753 and secured 1*^st^* place in the leaderboard. Not so impressively, we secured the 6*^th^* spot in the Lipo dataset leaderboard.

#### 3.1.2 Distribution benchmarks

The *distribution benchmarks* measure how a drug moves to and from the various tissues of the body and the amount of drug concentration in those tissues. These benchmarks consist of three datasets, two involving regression and one classification. The results of this benchmark were less satisfactory, particularly in the case of the classification dataset that measures the drug’s ability to penetrate the blood-brain barrier, ranking in the lower half of the entire benchmark. Notably, this particular benchmark appears to be among the most challenging, with only ensemble-based strategies showing the best performance. The results for the three benchmarks and respective pre-processing steps and models can be seen in Table 2.

**Table 2.**
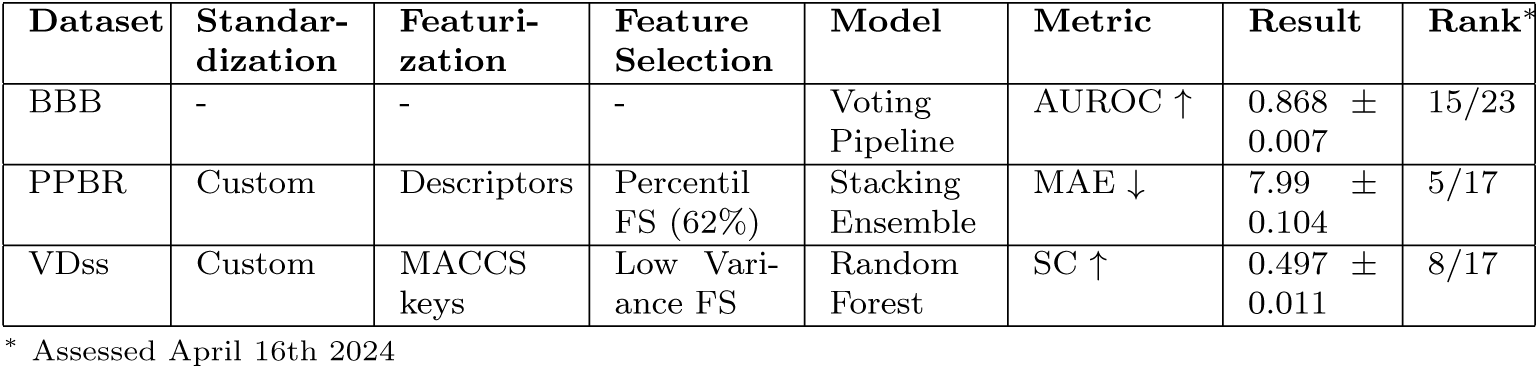
Performance of DeepMol in the three distribution-related properties in the TDC ADMET benchmark group.

#### 3.1.3 Metabolism benchmarks

The *metabolism benchmarks* evaluate the breakdown of drugs by specialized enzymatic systems, determining the duration and intensity of their effects. These benchmarks consist of six classification datasets. We achieved top-2 performance in two datasets and top-5 in three. Notably, unlike the absorption benchmarks, which thrived on graph-based approaches, simpler methods such as molecular descriptors and finger-prints yielded superior results in the metabolism benchmark. Similarly, ensemble learning techniques outperformed individual models in the *distribution benchmark*. The results for the six benchmarks and respective pre-processing steps and models can be seen in Table 3.

**Table 3.**
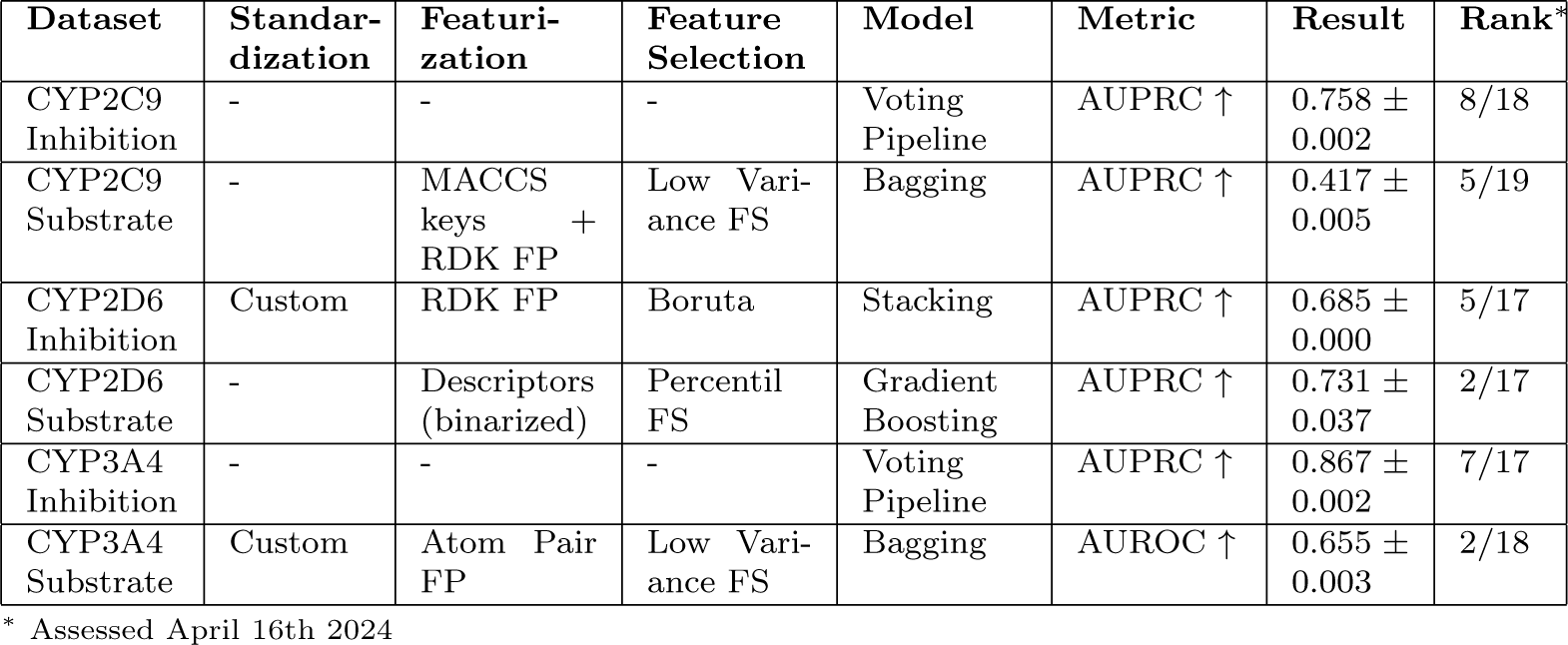
Performance of DeepMol in the six metabolism-related properties in the TDC ADMET benchmark group.

#### 3.1.4 Excretion benchmarks

The excretion benchmarks measure the removal of drugs from the body using various routes. These benchmarks consist of three regression datasets.

The exploration of predictive models across these datasets reveals that CNNs, alongside the ChEMBL standardizer, emerge as standout methodologies. For the Half-Life dataset, a DeepMol custom standardization and scaled descriptors combined with a 1D CNN achieved an SC score of 0.485, ranking it fifth. On the other hand, a voting pipeline that standardized molecules with the ChEMBL standardizer comprises three Directed Message Passing Neural Networks (D-MPNN) and two GCNs and reaches fourth place for the CL-Hepa dataset. For the CL-Micro dataset, optimal performance was achieved using a voting pipeline that combined five pipelines employing the ChEMBL standardizer with five distinct TextCNNs. The pipeline achieved a modest 11th place, lagging behind the first place by a small SC margin of 0.05.

The results for the three benchmarks and respective pre-processing steps and models can be seen in Table 4.

**Table 4.**
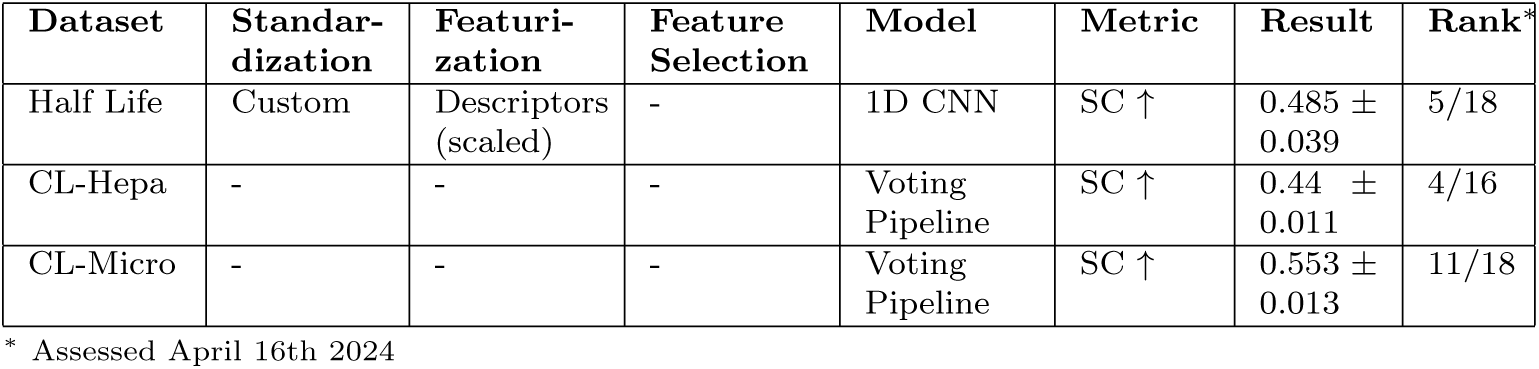
Performance of DeepMol in the three excretion-related properties in the TDC ADMET benchmark group.

#### 3.1.5 Toxicity benchmarks

The toxicity benchmarks measure how much damage a drug can cause to organisms. These benchmarks consist of four datasets, one involving regression and three for classification.

DeepMol achieved good results for the Ames and LD50 datasets, which ranked fifth and fourth, respectively. For the former dataset, a voting pipeline with five GNNs standardizing molecules with the ChEMBL standardized, while for the latter, a pipeline using a custom standardizer, layered FPs, an FS method that selected features based on the percentile 79 of the scores of a univariate linear regression test (for more information refer to [18]), and a voting regressor composed of 5 different models. On the other hand, the results for the rest of the datasets were far from impressive, with a GCN ending up in the 9th place of the hERG dataset leaderboard and a Voting pipeline securing a place in the half of the leaderboard for the DILI dataset.

The results for the four benchmarks and respective pre-processing steps and models can be seen in Table 5.

**Table 5.**
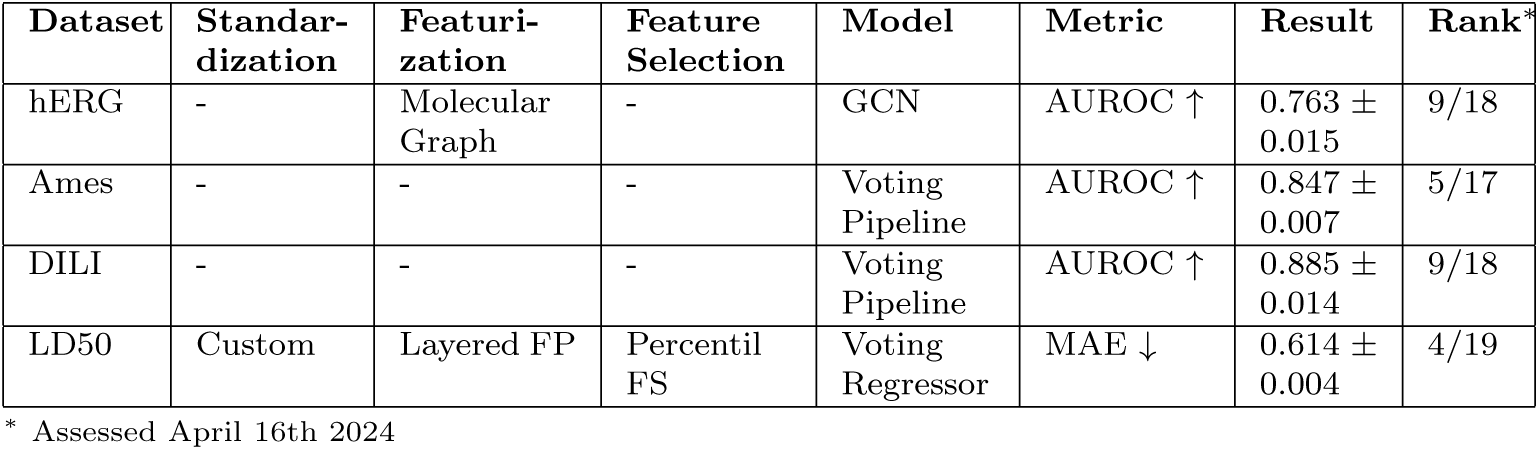
Performance of DeepMol in the four toxicity-related properties in the TDC ADMET benchmark group.

### 3.2 Practical applications of DeepMol

Several publications have already showcased the versatility of *DeepMol* in various domains, such as drug discovery, food science and natural product discovery, across different tasks. Numerous experiments have exploited the diverse array of methods offered by DeepMol, highlighting its user-friendly nature and contributing to significant research advancements.

In their studies [41, 53], Baptista and colleagues used *DeepMol* to investigate which compound representation methods are most suitable for drug sensitivity prediction in cancer cell lines. They benchmarked twelve compound representations, including molecular fingerprints and DL-based representation learning methods, using two classifications and three regression datasets from human cancer single-cell line drug screenings. The authors found that most compound representations performed similarly. Still, some end-to-end deep learning models performed on par with or even outperformed traditional fingerprint-based models, even when dealing with smaller datasets. Furthermore, the authors utilized *DeepMol* ’s feature importance methods to enhance the interpretability of fingerprint-based deep learning models. They demonstrated that consistently highlighted features were known to be associated with drug response.

Capela *et al.* [42] conducted a study in which they trained and evaluated 66 different model configurations to predict the relationships between chemical structures and sweetness. Throughout the study pipeline, *DeepMol* was utilized for several tasks, such as molecular standardization, feature generation, feature selection, model construction, hyperparameter tuning, and model explainability. Furthermore, a subset of the trained models was employed to screen 60 million molecules from PubChem [54] in search of potential sweeteners. The authors successfully identified numerous derivatives of potent and artificial sweeteners, some of which were patented as sweetening agents and were not included in the original training data. This demonstrated the remarkable capability of *DeepMol* in helping design new sweeteners and repurposing existing compounds.

In a recent study [55], DeepMol AutoML was used to address the challenge of predicting precursors of specialized plant metabolites, which play critical roles in plant defence and have significant economic implications. Despite these compounds’ complexity, vast diversity, and the current gaps in understanding their biosynthesis, DeepMol’s methodology stood out for its efficacy. It was useful to find regularized linear classifiers that surpass existing state-of-the-art approaches in performance and also offer chemical explanations for their predictions. This approach marks a significant advancement in expediting the discovery of biosynthetic pathways, highlighting the potential of DeepMol’s AutoML in finding high-performing models.

## 4 Discussion: comparison with other tools

Several chemoinformatics toolkits have been developed in recent years [56, 57, 58, 59, 10], including *DeepChem* (https://deepchem.io/), OpenChem [58], AMPL [57], and ZairaChem [16]. These open-source projects, developed in Python, are used to construct pipelines for QSAR/QSPR modeling. A comparison between DeepMol and the other tools is showcased in Figure 4.

**Fig. 4.**
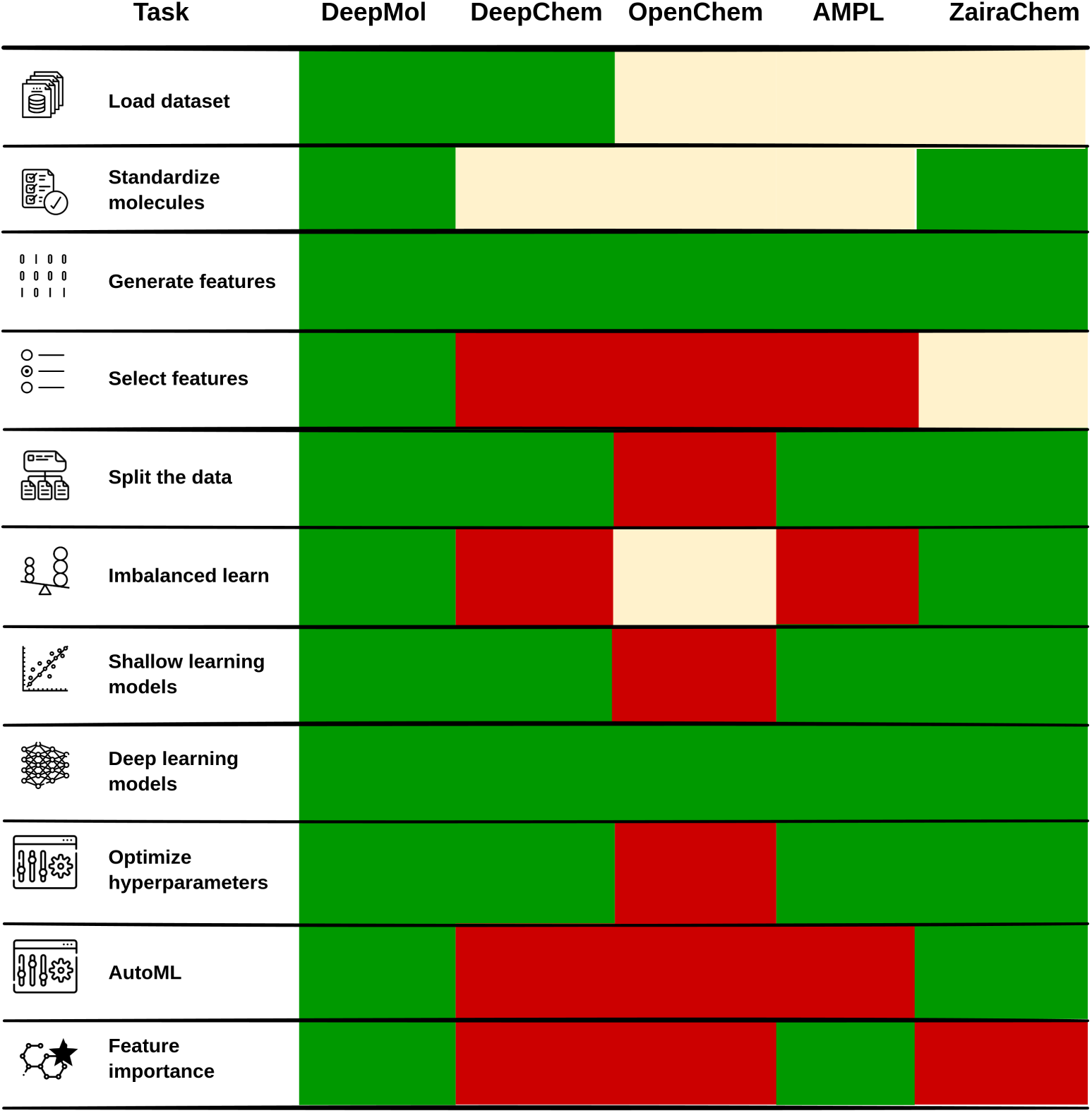
Heatmap showcasing the presence or absence of an integrated way of performing relevant tasks for constructing QSAR/QSPR systems. Green stands for presence, red for absence and yellow for incomplete integration.

Regarding dataset loading, *DeepMol* and *DeepChem* support CSV and SDF formats, while OpenChem, AMPL, and ZairaChem only support CSV files. Hence, a limitation of the latter two tools is their incapability to import pre-computed 3D structures. Molecule standardization is available in *DeepMol* through various methods. OpenChem allows sanitizing molecules, padding sequences and canonicalizing SMILES. In contrast, AMPL allows the striping of salt groups, neutralizing and kekulizing the molecules and replacing any rare isotopes with the most common ones for each element. *DeepChem* only allows sanitizing molecules when loading datasets. ZairaChem applies the MELLODDY-Tuner protocol, which includes disconnecting metal atoms from the rest of the molecule, reionizing the molecule, adjusting charges and protonation states, assigning stereochemistry to the molecule, ensuring consistent representation of chiral centres, and additional standardization procedures. All the tools can perform data splitting, except OpenChem, and only *DeepMol* can perform a stratified split specific to molecular data and multitask classification. Additionally, all the tools are capable of generating numerical features from molecules. Hyperparameter and architecture optimization are possible in all the toolkits except for OpenChem.

Model construction in these toolkits primarily focuses on DL models, but all the tools provide shallow learning models except for OpenChem. Hyperparameter and architecture optimization are possible in all the tools except OpenChem. More-over, while all the methods offer pipelines, only *DeepMol* and ZairaChem provide a comprehensive and efficient approach to automating pipeline optimization (AutoML). *DeepMol* excels compared to other tools in providing feature importance analysis and effectively connecting features to molecular structures. Both ZairaChem and *DeepMol* also offer methods to address unbalanced datasets and perform integrated and automated feature selection, distinguishing them from the other assessed tools. ZairaChem applies feature selection through Autogluon, which offers only one feature selection method.

Even though *DeepMol* and ZairaChem appear to perform similarly for these tasks, it is important to note that ZairaChem is limited to binary classification problems. Consequently, the features provided by ZairaChem, as indicated in Figure 4, are specifically designed for binary classification tasks.

## 5 Conclusion

In conclusion, *DeepMol* emerges as a powerful alternative to similar tools, offering a Python-based open-source framework for predicting activities and properties of chemical molecules. Its modular design allows researchers to customize every aspect of the ML pipeline, from data processing to model prediction and explainability, catering to users with varying levels of computational expertise. *DeepMol* ’s AutoML modules further simplify the process by automatically optimizing pre-processing, data engineering techniques, and ML/DL models and their hyperparameters, streamlining the selection of the most suitable combinations for a given dataset. By providing a user-friendly and easily customizable platform, *DeepMol* empowers researchers to efficiently explore thousands of configurations, making it an invaluable resource for accelerating chemical discovery. With the support of well-established packages like *RDKit*, *Scikit-Learn*, *Tensorflow*, and *DeepChem*, *DeepMol* represents a promising avenue for optimizing and deploying pipelines, offering a vital contribution to the advancement of computational chemistry. The framework’s availability as an open-source resource encourages collaboration and innovation in the field, facilitating progress and empowering researchers in their quest for finding new chemicals with improved properties.

## Acknowledgements

This study was supported by the Portuguese Foundation for Science and Technology(FCT) under the scope of the strategic funding of UIDB/04469/2020 unit, and by LABBELS – Associate Laboratory in Biotechnology, Bioengineering and Microelectromechanical Systems, LA/P/0029/2020. Moreover, this research has been supported by the Portuguese Foundation for Science and Technology (FCT) through the DeepBio project - ref. NORTE-01-0247-FEDER-039831, funded by Lisboa 2020, Norte 2020, Portugal 2020 and the project SHIKIFACTORY100 - Modular cell factories for the production of 100 compounds from the shikimate pathway (Reference 814408). We also thank FCT for the PhD fellowships to J. Capela (DFA/BD/08789/2021) and J. Correia (SFRH/BD/144314/2019).

## Notes

https://github.com/BioSystemsUM/DeepMol

### Competing Interest Statement

The authors have declared no competing interest.

## References

[1] Gerhard Hessler and Karl-Heinz Baringhaus. “Artificial Intelligence in Drug Design”. In: Molecules 23.10 (Oct. 2018), p. 2520. doi: 10.3390/molecules23102520. url: https://doi.org/10.3390/molecules23102520.

[2] Jie Shen and Christos A. Nicolaou. “Molecular property prediction: recent trends in the era of artificial intelligence”. In: Drug Discovery Today: Technologies 32–33 (Dec. 2019), pp. 29–36. doi: 10.1016/j.ddtec.2020.05.001. url: https://doi.org/10.1016/j.ddtec.2020.05.001.

[3] Johann Gasteiger. “Chemistry in Times of Artificial Intelligence”. In: ChemPhysChem 21.20 (Sept. 2020), pp. 2233–2242. doi: 10.1002/cphc.202000518. url: https://doi.org/10.1002/cphc.202000518.

[4] W. Patrick Walters and Regina Barzilay. “Applications of Deep Learning in Molecule Generation and Molecular Property Prediction”. In: Accounts of Chemical Research 54.2 (Dec. 2020), pp. 263–270. doi: 10.1021/acs.accounts.0c00699. url: https://doi.org/10.1021/acs.accounts.0c00699.

[5] Grégoire Montavon, et al. “Machine learning of molecular electronic properties in chemical compound space”. In: New Journal of Physics 15 (9 Sept. 2013), p. 095003. issn: 1367–2630. doi: 10.1088/1367-2630/15/9/095003.

[6] Alexandre Tkatchenko. “Machine learning for chemical discovery”. In: Nature Communications 11 (1 Aug. 2020), p. 4125. issn: 2041-1723. doi: 10.1038/s41467-020-17844-8.

[7] Mati Karelson, Victor S. Lobanov, and Alan R. Katritzky. “Quantum-Chemical Descriptors in QSAR/QSPR Studies”. In: Chemical Reviews 96 (3 Jan. 1996), pp. 1027–1044. issn: 0009-2665. doi: 10.1021/cr950202r.

[8] Workalemahu M. Berhanu et al. “Quantitative Structure-Activity/Property Relationships: The Ubiquitous Links between Cause and Effect”. In: ChemPlusChem 77 (7 July 2012), pp. 507–517. issn: 21926506. doi: 10.1002/cplu.201200038.

[9] Paulo C. S. Costa et al. “Chemical Graph Theory for Property Modeling in QSAR and QSPR—Charming QSAR QSPR”. In: Mathematics 9 (1 Dec. 2020), p. 60. issn: 2227-7390. doi: 10.3390/math9010060.

[10] Isadora Leitzke Guidotti et al. “Bambu and its applications in the discovery of active molecules against melanoma”. In: Journal of Molecular Graphics and Modelling 124 (Nov. 2023), p. 108564. issn: 10933263. doi: 10.1016/j.jmgm.2023.108564.

[11] Gregory Sliwoski et al. “Computational Methods in Drug Discovery”. In: Pharmacological Reviews 66.1 (2014). Ed. by Eric L. Barker, pp. 334–395. issn: 0031-6997. doi: 10.1124/pr.112.007336. eprint: https://pharmrev.aspetjournals.org/content/66/1/334.full.pdf. url: https://pharmrev.aspetjournals.org/content/66/1/334.

[12] Zhenxing Wu et al. “Do we need different machine learning algorithms for QSAR modeling? A comprehensive assessment of 16 machine learning algorithms on 14 QSAR data sets”. In: Briefings in Bioinformatics 22 (4 July 2021). issn: 1467-5463. doi: 10.1093/bib/bbaa321.

[13] Yuting Xu. “Deep Neural Networks for QSAR”. In: 2022, pp. 233–260. doi: 10.1007/978-1-0716-1787-810.

[14] Zhen Li et al. “Deep learning methods for molecular representation and property prediction”. In: Drug Discovery Today 27.12 (Dec. 2022), p. 103373. doi: 10.1016/j.drudis.2022.103373. url: https://doi.org/10.1016/j.drudis.2022.103373.

[15] Álmos Orosz, Károly Héberger, and Anita Rácz. “Comparison of Descriptor- and Fingerprint Sets in Machine Learning Models for ADME-Tox Targets”. In: Frontiers in Chemistry 10 (June 2022). doi: 10.3389/fchem.2022.852893. url: https://doi.org/10.3389/fchem.2022.852893.

[16] G. Turon et al. “First fully-automated AI/ML virtual screening cascade implemented at a drug discovery centre in Africa”. In: Nat Commun 14 (2023), p. 5736. doi: 10.1038/s41467-023-41512-2. url: https://doi.org/10.1038/s41467-023-41512-2.

[17] RDKit: Open-source cheminformatics. http://www.rdkit.org.

[18] F. Pedregosa et al. “Scikit-learn: Machine Learning in Python”. In: Journal of Machine Learning Research 12 (2011), pp. 2825–2830.

[19] Martín Abadi et al. TensorFlow: Large-Scale Machine Learning on Heterogeneous Systems. Software available from tensorflow.org. 2015. url: https://www.tensorflow.org/.

[20] Bharath Ramsundar et al. Deep Learning for the Life Sciences. https://www.amazon.com/Deep-Learning-Life-Sciences-Microscopy/dp/1492039837. O’Reilly Media, 2019.

[21] Takuya Akiba et al. “Optuna: A Next-generation Hyperparameter Optimization Framework”. In: Proceedings of the 25th ACM SIGKDD International Conference on Knowledge Discovery and Data Mining. 2019.

[22] Kexin Huang et al. “Therapeutics Data Commons: Machine Learning Datasets and Tasks for Drug Discovery and Development”. In: Proceedings of the Neural Information Processing Systems Track on Datasets and Benchmarks. Ed. by J. Vanschoren and S. Yeung. Vol. 1. Curran, 2021. url: https://datasets-benchmarks-proceedings.neurips.cc/paper_files/paper/2021/file/4c56ff4ce4aaf9573aa5dff913df997a-Paper-round1.pdf.

[23] Daniel Probst and Jean-Louis Reymond. “SmilesDrawer: Parsing and Drawing SMILES-Encoded Molecular Structures Using Client-Side JavaScript”. In: Journal of Chemical Information and Modeling 58.1 (2018). PMID: 29257869, pp. 1–7. doi: 10.1021/acs.jcim.7b00425. eprint: https://doi.org/10.1021/acs.jcim.7b00425. url: https://doi.org/10.1021/acs.jcim.7b00425.

[24] Johann Gasteiger et al. “Chemical Information in 3D Space”. In: Journal of Chemical Information and Computer Sciences 36.5 (1996), pp. 1030–1037. doi: 10.1021/ci960343+. eprint: https://doi.org/10.1021/ci960343+. url: https://doi.org/10.1021/ci960343+.

[25] Jaroslaw Polanski and Johann Gasteiger. “Computer representation of chemical compounds”. In: Handbook of Computational Chemistry. Springer International Publishing, Jan. 2017, pp. 1997–2039. isbn: 9783319272825. doi: 10.1007/978-3-319-27282-550.

[26] Daniel S. Wigh, Jonathan M. Goodman, and Alexei A. Lapkin. “A review of molecular representation in the age of machine learning”. In: WIREs Computational Molecular Science 12 (5 Sept. 2022). issn: 1759-0876. doi: 10.1002/wcms.1603.

[27] K. Mansouri et al. “An automated curation procedure for addressing chemical errors and inconsistencies in public datasets used in QSAR modelling”. In: SAR and QSAR in Environmental Research 27 (11 Nov. 2016), pp. 911–937. issn: 1062-936X. doi: 10.1080/1062936X.2016.1253611.

[28] Denis Fourches, Eugene Muratov, and Alexander Tropsha. “Trust, But Verify: On the Importance of Chemical Structure Curation in Cheminformatics and QSAR Modeling Research”. In: Journal of Chemical Information and Modeling 50.7 (June 2010), pp. 1189–1204. doi: 10.1021/ci100176x. url: https://doi.org/10.1021/ci100176x.

[29] A. Patrícia Bento, et al. “An open source chemical structure curation pipeline using RDKit”. In: Journal of Cheminformatics 12.1 (Sept. 2020). doi: 10.1186/s13321-020-00456-1. url: https://doi.org/10.1186/s13321-020-00456-1.

[30] Volker D. Hähnke, Sunghwan Kim, and Evan E. Bolton. “PubChem chemical structure standardization”. In: Journal of Cheminformatics 10.1 (Aug. 2018). doi: 10.1186/s13321-018-0293-8. url: https://doi.org/10.1186/s13321-018-0293-8.

[31] Karen Karapetyan et al. “The Chemical Validation and Standardization Platform (CVSP): large-scale automated validation of chemical structure datasets”. In: Journal of Cheminformatics 7.1 (June 2015). doi: 10.1186/s13321-015-0072-8. url: https://doi.org/10.1186/s13321-015-0072-8.

[32] Viviana Consonni and Roberto Todeschini. Molecular descriptors for chemoinformatics. John Wiley & Sons, 2009.

[33] Roberto Todeschini and Viviana Consonni. “Molecular descriptors”. In: Recent Advances in QSAR Studies (2010), pp. 29–102.

[34] Hirotomo Moriwaki et al. “Mordred: a molecular descriptor calculator”. In: Journal of Cheminformatics 10 (1 Dec. 2018), p. 4. issn: 1758-2946. doi: 10.1186/s13321-018-0258-y.

[35] Xinhao Li and Denis Fourches. “SMILES Pair Encoding: A Data-Driven Substructure Tokenization Algorithm for Deep Learning”. In: Journal of Chemical Information and Modeling 61 (4 Apr. 2021), pp. 1560–1569. issn: 1549-9596. doi: 10.1021/acs.jcim.0c01127.

[36] Steven Kearnes et al. “Molecular graph convolutions: moving beyond finger-prints”. In: Journal of Computer-Aided Molecular Design 30 (8 Aug. 2016), pp. 595–608. issn: 0920-654X. doi: 10.1007/s10822-016-9938-8.

[37] David K Duvenaud, et al. “Convolutional networks on graphs for learning molecular fingerprints”. In: Advances in neural information processing systems 28 (2015).

[38] Nicola De Cao and Thomas Kipf. “MolGAN: An implicit generative model for small molecular graphs”. In: arXiv preprint arXiv:1805.11973 (2018).

[39] Grégoire Montavon, et al. “Learning invariant representations of molecules for atomization energy prediction”. In: Advances in neural information processing systems 25 (2012).

[40] Miron B. Kursa and Witold R. Rudnicki. “Feature Selection with thebBoruta/bPackage”. In: Journal of Statistical Software 36.11 (2010). doi: 10.18637/jss.v036.i11. url: https://doi.org/10.18637/jss.v036.i11.

[41] Delora Baptista et al. “Evaluating molecular representations in machine learning models for drug response prediction and interpretability”. In: Journal of Integrative Bioinformatics 19.3 (Aug. 2022). doi: 10.1515/jib-2022-0006. url: https://doi.org/10.1515/jib-2022-0006.

[42] Joao Capela et al. “Development of Deep Learning approaches to predict relationships between chemical structures and sweetness”. In: 2022 International Joint Conference on Neural Networks (IJCNN). IEEE, July 2022. doi: 10.1109/ijcnn55064. 2022. 9891992. url: https://doi.org/10.1109/ijcnn55064.2022.9891992.

[43] Darko Butina. “Unsupervised Data Base Clustering Based on Daylight’s Finger-print and Tanimoto Similarity: A Fast and Automated Way To Cluster Small and Large Data Sets”. In: Journal of Chemical Information and Computer Sciences 39.4 (1999), pp. 747–750. doi: 10.1021/ci9803381. eprint: https://doi.org/10.1021/ci9803381. url: https://doi.org/10.1021/ci9803381.

[44] Congying Xu et al. “In silico prediction of chemical Ames mutagenicity”. In: Journal of chemical information and modeling 52.11 (2012), pp. 2840–2847.

[45] Justin M. Johnson and Taghi M. Khoshgoftaar. “Survey on deep learning with class imbalance”. In: Journal of Big Data 6.1 (Mar. 2019). doi: 10.1186/s40537-019-0192-5. url: https://doi.org/10.1186/s40537-019-0192-5.

[46] Seļcuk Korkmaz. “Deep Learning-Based Imbalanced Data Classification for Drug Discovery”. In: Journal of Chemical Information and Modeling 60.9 (June 2020), pp. 4180–4190. doi: 10.1021/acs.jcim.9b01162. url: https://doi.org/10.1021/acs.jcim.9b01162.

[47] Guillaume Lemaître, Fernando Nogueira, and Christos K. Aridas. “Imbalanced-learn: A Python Toolbox to Tackle the Curse of Imbalanced Datasets in Machine Learning”. In: Journal of Machine Learning Research 18.17 (2017), pp. 1–5. url: http://jmlr.org/papers/v18/16-365.html.

[48] N. V. Chawla et al. “SMOTE: Synthetic Minority Over-sampling Technique”. In: Journal of Artificial Intelligence Research 16 (June 2002), pp. 321–357. doi: 10.1613/jair.953. url: https://doi.org/10.1613/jair.953.

[49] Gustavo E. A. P. A. Batista, Ana Lúcia Cetertich Bazzan, and Maria Carolina Monard. “Balancing Training Data for Automated Annotation of Keywords: a Case Study”. In: WOB. 2003.

[50] Gustavo E. A. P. A. Batista, Ronaldo C. Prati, and Maria Carolina Monard. “A study of the behavior of several methods for balancing machine learning training data”. In: ACM SIGKDD Explorations Newsletter 6.1 (June 2004), pp. 20–29. doi: 10.1145/1007730.1007735. url: https://doi.org/10.1145/1007730.1007735.

[51] Scott M Lundberg and Su-In Lee. “A unified approach to interpreting model predictions”. In: Advances in neural information processing systems 30 (2017).

[52] Chang-Ying Ma et al. “Prediction models of human plasma protein binding rate and oral bioavailability derived by using GA–CG–SVM method”. In: Journal of pharmaceutical and biomedical analysis 47.4–5 (2008), pp. 677–682.

[53] Delora Baptista et al. “A Comparison of Different Compound Representations for Drug Sensitivity Prediction”. In: Practical Applications of Computational Biology &amp Bioinformatics, 15th International Conference (PACBB 2021). Springer International Publishing, Aug. 2021, pp. 145–154. doi: 10.1007/978-3-030-86258-915. url: https://doi.org/10.1007/978-3-030-86258-915.

[54] Sunghwan Kim, et al. “PubChem 2023 update”. In: Nucleic Acids Research 51.D1 (Oct. 2022), pp. D1373–D1380. doi: 10.1093/nar/gkac956. url: https://doi.org/10.1093/nar/gkac956.

[55] João Capela, et al. “Automated Machine Learning to Predict the Precursors of Plant Specialized Metabolites”. In: Manuscript submitted (2024).

[56] Sujit R. Tangadpalliwar et al. “ChemSuite: A package for chemoinformatics calculations and machine learning”. In: Chemical Biology Drug Design 93.5 (May 2019), pp. 960–964. issn: 1747-0277. doi: 10.1111/cbdd.13479. url: https://onlinelibrary.wiley.com/doi/10.1111/cbdd.13479.

[57] Amanda J. Minnich et al. “AMPL: A Data-Driven Modeling Pipeline for Drug Discovery”. In: Journal of Chemical Information and Modeling 60.4 (Apr. 2020), pp. 1955–1968. issn: 1549-9596. doi: 10.1021/acs.jcim.9b01053. url: https://pubs.acs.org/doi/10.1021/acs.jcim.9b01053.

[58] Maria Korshunova et al. “OpenChem: A Deep Learning Toolkit for Computational Chemistry and Drug Design”. In: Journal of Chemical Information and Modeling 61.1 (Jan. 2021), pp. 7–13. issn: 1549-9596. doi: 10.1021/acs.jcim.0c00971. url: https://pubs.acs.org/doi/10.1021/acs.jcim.0c00971.

[59] Benjamin P. Brown et al. “Introduction to the BioChemical Library (BCL): An Application-Based Open-Source Toolkit for Integrated Cheminformatics and Machine Learning in Computer-Aided Drug Discovery”. In: Frontiers in Pharmacology 13 (Feb. 2022). issn: 1663-9812. doi: 10.3389/fphar.2022.833099. url: https://www.frontiersin.org/articles/10.3389/fphar.2022.833099/full.

