## Supplementary Table for "DeepMol: An Automated Machine and Deep Learning Framework for Computational Chemistry"

### Extract features

In machine learning, extracting features from molecules is a common task. Molecular features can be categorized into four types: 0D, 1D, 2D, 3D, and 4D. 0D features provide information about the entire molecule, including properties like atom count, bond count, and molecular weight. 1D features describe substructures within the molecule, such as molecular fingerprints and fragment keys. 2D features capture the molecular topology based on the graph representation, including the number of rings and rotatable bonds. 3D features capture the geometric descriptors of the molecule's three-dimensional structure. Lastly, 4D features introduce an additional dimension to capture interactions between the molecule and an active site or multiple conformational states, such as molecular dynamics.

As we move from 0D to 4D molecular descriptors, the computational cost of feature calculation increases. For instance, generating 3D features involves creating 3D conformers, which can be time-consuming for larger molecules. It's worth noting that certain features may not be applicable to all molecules; for example, 3D features cannot be calculated for molecules lacking a 3D structure.

DeepMol provides a wide set of 1D features, all provided by rdkit. DeepMol includes one of the most famous circular fingerprints, the Extended Connectivity Fingerprint (ECFP), atom pair fingerprints that encode the presence or absence of pairs of atoms in a molecule, as well as the distance between them. Moreover, DeepMol includes layered fingerprints that find all possible paths or subgraphs of specified lengths in the molecule based on the input parameters and compute layers of structural and functional features per molecular subgraph. DeepMol also includes an RDKit-specific fingerprint inspired by public descriptions of the Daylight fingerprint. The fingerprinting algorithm generates molecular fingerprints by identifying subgraphs within a specified size range, hashing each subgraph to create a raw bit that is hashed to fit the fingerprint size. The default scheme for subgraph hashing involves considering factors such as atom types (based on atomic number and aromaticity), atom degrees in the path, and bond types. Finally, DeepMol includes Molecular ACCess System (MACCS) keys that encode the presence or absence of certain molecular fragments or substructures in a molecule as a binary bitstring. The fragments used are based on a predefined set of SMARTS patterns, which represent specific substructures or features of a molecule.

DeepMol also provides a set of 0D, a few 1D and 2D descriptors in only one class. Those are enumerated and described in Table S1 and at https://deepmol.readthedocs.io/en/latest/deepmol_docs/featurization.html#d-1d-and-2d-descriptors.

As mentioned above, 3D molecular conformations have to be generated or loaded prior to generating 3D descriptors. For this matter, the conformer generation process begins by utilizing the Experimental-Torsion basic Knowledge Distance Geometry (ETKDG) algorithm, which is an extension of the Knowledge Distance Geometry (KDG) method. ETKDG incorporates efficiency enhancements and knowledge-based rules to generate a diverse set of low-energy conformers for small organic molecules. This method combines random sampling and efficient energy evaluations to strike a balance between computational efficiency and conformational coverage. It is widely employed in molecular modelling and drug discovery applications. Subsequently, the Merck Molecular Force Field (MMFF) and Universal Force Field (UFF) algorithms are employed. These force fields optimize the conformers by calculating the potential energy and atomic forces based on the molecule's geometry. This process guides the conformational search towards more stable conformations.

Once generated, the methods within DeepMol can be used to extract features from the conformers. These methods encompass \textit{AutoCorr3D} that captures spatial autocorrelation patterns in a molecule, while the Radial Distribution Function (RDF) characterizes the distribution of particles based on their distances from a reference particle. The plane of best fit determines the optimal plane that fits a set of points, and MORSE utilizes molecular transforms to derive information from atomic coordinates. WHIM descriptors provide a holistic representation of a molecule's structure, while the Radius of Gyration measures its spatial extent. The Inertial Shape Factor, Eccentricity, Asphericity, and Spherocity Index quantify the shape and symmetry of the molecule. Principal Moments of Inertia describe its rotational behaviour, and Normalized Principal Moments Ratios provide insight into the relative magnitudes of the principal moments. These descriptors collectively contribute to a comprehensive understanding of a molecule's three-dimensional properties and structural characteristics.

Table S1 - 0D, 1D and 2D descriptors integrated into only one class in Descriptors

| Set of descriptors | Descriptors | Description |
|---|---|---|
| EState index descriptors | MaxAbsEStateIndex | Maximum absolute EState index - The MAEstate specifically represents the highest absolute EState index value among all the atoms in a molecule. It indicates the atom with the largest charge magnitude, reflecting its potential reactivity or contribution to chemical properties. |
| | MaxEStateIndex | Maximum EState Index - The MaxEStateIndex specifically represents the highest EState index value among all the atoms in a molecule. It indicates the atom with the largest charge or electronic density, reflecting its potential reactivity or significance in the molecule’s properties. |
| | MinAbsEStateIndex | Minimum absolute EState index - The MinAbsEStateIndex specifically represents the lowest absolute EState index value among all the atoms in a molecule. |
| | MinEStateIndex | Minimum EState index - The MinEStateIndex represents the lowest EState index value among all the atoms in a molecule. |
| QED | QED | Quantitative estimation of drug-likeness - a computational algorithm used to quantitatively assess the drug-likeness of a molecule. It combines various molecular descriptors, including 2D properties, to generate a single numerical score that represents the overall drug-likeness of the molecule. |
| Molecular weight descriptors | MolWt | Molecular weight. |
| | HeavyAtomMolWt | The average molecular weight of the molecule, ignoring hydrogens. |
| | ExactMolWt | The exact molecular weight of the molecule. |
| Electron descriptors | NumValenceElectrons | The number of valence electrons the molecule has. |
| | NumRadicalElectrons | The number of radical electrons the molecule has. |
| Charge descriptors | MaxPartialCharge | Maximum partial charge |
| | MinPartialCharge | Minimum partial charge |

| | | |
|---|---|---|
| | MaxAbsPartialCharge | Maximum absolute partial charge |
| | MinAbsPartialCharge | Minimum absolute partial charge |
| Morgan fingerprint density | FpDensityMorgan1 | Quantify the frequency of occurrence of specific substructures within the molecule at a local level, taking into account their immediate surroundings. Higher values of density indicate a higher density of unique substructures in the molecule, while lower values indicate fewer unique substructures or a more uniform distribution of substructures. Densities for Morgan radius 1, 2 and 3. |
| | FpDensityMorgan2 | |
| | FpDensityMorgan3 | |
| BCUT2D descriptors | BCUT2D_MWHI | Incorporates atom masses in the Burden matrix - returns the highest eigenvalue. |
| | BCUT2D_MWLOW | Incorporates atom masses in the Burden matrix - returns the lowest eigenvalue. |
| | BCUT2D_CHGHI | Incorporates atom charges in the Burden matrix - returns the highest eigenvalue. |
| | BCUT2D_CHGLO | Incorporates atom charges in the Burden matrix - returns the lowest eigenvalue. |
| | BCUT2D_LOGPHI | Incorporates atom logarithms of the partition coefficient (logP) in the Burden matrix - returns the highest eigenvalue. |
| | BCUT2D_LOGPLOW | Incorporates atom logarithms of the partition coefficient (logP) in the Burden matrix - returns the lowest eigenvalue. |
| | BCUT2D_MRHI | Incorporates atom molar refractivity in the Burden matrix - returns the highest eigenvalue. |
| | BCUT2D_MRLOW | Incorporates atom molar refractivity in the Burden matrix - returns the lowest eigenvalue. |
| AvgIpc | AvgIpc | The average information content of the coefficients of the characteristic polynomial of the adjacency matrix of a hydrogen-suppressed graph of a molecule. |
| Ipc | Ipc | The information content of the coefficients of the characteristic polynomial of the adjacency matrix of a hydrogen-suppressed graph of a molecule. |

| | | |
|---|---|---|
| BalabanJ | BalabanJ | Balaban's J index. It quantifies the molecular topological structure by considering the connectivity of atoms and bonds in the molecule. |
| BertzCT | BertzCT | Bertz complexity index. It measures the topological complexity or branching of a molecule based on its structural connectivity. |
| Chi descriptors | Chi descriptors | The Chi descriptors represent the count of specific path patterns in the molecule and are calculated based on the Hall-Kier delta values or on the deviation of an atom's valence electron count from the expected count based on its atomic number. |
| HallKierAlpha | HallKierAlpha | Hall-Kier alpha value. It describes the flexibility or rigidity of atoms in a molecule. |
| Kappa descriptors | Kappa1 | Kappa shape indices. They describe the shape of a molecule based on the distribution of bond lengths and angles. These descriptors are derived from the Hall-Kier alpha descriptor and the number of paths of specific lengths in the molecule. |
| | Kappa2 | |
| | Kappa3 | |
| LabuteASA | LabuteASA | Labute's Approximate Surface Area. It estimates the solvent-accessible surface area of a molecule, which is relevant for its solubility and permeability properties. |
| PEOE VSA descriptors | PEOE VSA descriptors | These descriptors Calculates the PEOE (Partial Equalization of Orbital Electronegativity) VSA (surface area contributions of atoms or groups of atoms in a molecule) for a molecule by assigning atom contributions to predefined bins based on their Labute ASA and Gasteiger charge values. |
| SMR VSA descriptors | SMR VSA descriptors | Calculates the SMR (Molar Refractivity) VSA for a molecule by assigning atom contributions to predefined bins based on their Labute ASA and MR values. |
| SlogP VSA descriptors | SlogP VSA descriptors | Calculates the SlogP VSA for a molecule by assigning atom contributions to predefined bins based on their Labute ASA and SlogP values. |
| EState VSA | EState VSA | Calculates the EState (E-State) VSA for a molecule by assigning atom contributions to predefined bins based on their Labute ASA and EState values. |
| FractionCSP3 | FractionCSP3 | Fraction of sp3-hybridized carbon atoms in the molecule. |

| | | |
|---|---|---|
| MolLogP | MolLogP | Molar logarithm of the partition coefficient (logP). It quantifies the lipophilicity or hydrophobicity of a molecule, which is important for its distribution and permeability properties. |
| TPSA | TPSA | Topological polar surface area. It estimates the surface area of a molecule that is involved in polar interactions, which is relevant for its solubility and biological activity. |
| RingCount | RingCount | Number of rings in the molecule. It indicates the level of molecular complexity and rigidity. |
| Count of structural groups | Count of structural groups | These descriptors count the number of hydrogen bond acceptor groups, hydrogen bond donor groups, heteroatoms (non-carbon atoms), etc., present in the molecule. They provide information about the potential for specific molecular interactions. |
| Frequency of functional groups | Frequency of functional groups | These descriptors represent the count of specific functional groups or substructures in the molecule. They provide information about the presence of particular chemical moieties. |

Table S2 - Molecular Fingerprints provided on DeepMol

| Molecular Fingerprint | Description | References |
|---|---|---|
| MorganFingerprint | Circular fingerprint based on the Morgan algorithm. This fingerprint is generated by considering the “circular” environment of each atom up to a given radius. | [1] |
| MACCSkeysFingerprint | Uses the 166 public keys implemented as SMARTS. The fragment definitions for the MACCS 166 keys can be found in this document: https://github.com/rdkit/rdkit/blob/master/rdkit/Chem/MACCSkeys.py | [2] |
| RDKFingerprint | This fingerprinting identifies all subgraphs in the molecule within a particular range of sizes and hashes each subgraph based on atom and bond types and degrees. Based on the Daylight fingerprint. | [3] |

| | | |
|---|---|---|
| LayeredFingerprint | Uses the same subgraph enumeration algorithm used in the RDKFingerprint. Sets bits based on different atom and bond type definitions (pure topology, bond order, atom types, aromaticity, etc). | - |
| AtomPairFingerprint | This fingerprint is generated by encoding interactions between all possible atom pairs in a molecule using a hashing scheme. These interactions are encoded based on atomic environments and shortest path between the atom pair. | [4] |

Table S3 - 3D descriptors provided on DeepMol

| 3D Descriptor | Description | Reference |
|---|---|---|
| AutoCorr3D | These descriptors are derived from the autocorrelation of various physicochemical properties such as charge, mass, van der Waals volume, electronegativity, polarizability, ionization potential, and electron affinity associated with the atoms within a molecule. In total, there are 80 descriptors. | [5] |
| RadialDistribution Function | These descriptors are based on the radial distribution function, describing the likelihood of finding an atom at a specific distance from another atom. They provide information about the spatial distribution of atoms and their environments. In total, there are 210 descriptors. | [5] |
| PlaneOfBestFit | The Plane of Best Fit (PBF) is the geometric plane that minimizes the sum of squared distances between the atoms of a molecule and the plane itself. This descriptor indicates the average distance of all heavy atoms from the PBF, offering a quantitative measure of how much the molecule deviates from a 2D shape and providing insight into its 3D configuration. | [6] |

| | | |
|---|---|---|
| MORSE | 224 Molecular Surface Electrostatics derive from the electrostatic potentials present on the molecular surface. They offer insights into the charge distribution across the molecule's surface and contribute to understanding its three-dimensional shape. In total, thera are 224 descriptors. | [5] |
| WHIM | Weighted Holistic Invariant Molecular (WHIM) descriptors are based on the principle of invariance, ensuring that they maintain consistency even after molecule transformations or rotations. They depend on statistical indexes obtained by projecting atoms along principal axes. WHIM descriptors encapsulate three-dimensional information regarding molecular size, shape, symmetry, and atom distribution, all in reference to invariant reference frames. In total, there are 114 descriptors. | [5] |
| RadiusOfGyration | The radius of gyration quantifies how atoms are distributed in a molecular structure cregarding its center of mass. Put simply, it represents the average distance of a molecule's atoms from their center of mass. | [7] |
| PrincipalMomentsOfInertia | The principal moments of inertia refer to the three rotational inertia values around its principal axes. These axes are the mutually perpendicular axes through the center of mass of the molecule, and the moments of inertia represent the distribution of mass around each axis. | - |
| InertialShapeFactor | The inertial shape factor is a measure that characterizes how mass is distributed around the principal axes of rotation. It is derived from the principal moments of inertia. | [5] |
| Eccentricity | The eccentricity refers to the extent to which its principal axes differ in length. It is calculated as the square root of the ratio of the difference between the squares of the longest and shortest principal moments of inertia to the square of the sum of all three principal moments of inertia. | [7] |

| | | |
|---|---|---|
| Asphericity | The asphericity of a molecule is a measure that quantifies the deviation of its shape from a perfect sphere. | [8] |
| SpherocityIndex | The spherosity index is an anisometry descriptor defined as a function of the eigenvalues of the covariance matrix of the atomic coordinates. It varies from zero for flat molecules, such as benzene, to unity for totally spherical molecules | [5] |
| NormalizedPrincipalMomentsRatios | Normalized ratios of principal moments of inertia measure the distribution of mass around the principal axes of rotation for a molecule, providing insights into its overall shape and symmetry. 2 descriptors. | [9] |

Table S4 - DeepChem featurizers provided by DeepMol

| Featurizer | Description | References |
|---|---|---|
| ConvMolFeat | Featurization to implement Duvenaud graph convolutions. It constructs a vector of local descriptors for each atom in a molecule. | [10] |
| PagtnMolGraphFeat | It creates a molecular graph connecting all atom pairs, considering interactions between every atom pair in the molecule. The default node representation includes features such as atom type, formal charge, degree, explicit and implicit valence, and aromaticity, resulting in a feature length of 94. The default edge representation, with a feature length of 42, considers bond type, conjugation, same ring membership, ring size, aromaticity, and distance between atom pairs based on the shortest path. | [11] |
| WeaveFeat | Featurization to implement Weave convolutions. In contrast to Duvenaud graph convolutions, Weave convolutions require a quadratic matrix of interaction | [12] |

| | | |
|---|---|---|
| | descriptors for every atom pair, potentially offering enhanced descriptive capability but resulting in a larger featurized dataset. | |
| MolGanFeat | This featurizer was originaly designed for MolGAN de-novo molecular generation. It encapsulates two matrices containing atom and bond type information that can be used in predictive models. | [13] |
| MolGraphConvFeat | This serves as a featurizer for general graph convolution networks applied to molecules, with default node and edge representations based on the WeaveNet paper. The default node features encompass atom type, formal charge, hybridization, hydrogen bonding, aromaticity, degree, number of hydrogens, chirality, and partial charge, while the default edge features include bond type, same ring membership, conjugation, and stereo configuration. Users have the flexibility to customize their own representations. | [12] |
| CoulombFeat | This featurizer calculates Coulomb matrices for molecules, offering a representation of the electronic structure. The resulting Coulomb matrix is an N x N matrix, where N represents the number of atoms in the molecule, with each element indicating the strength of the electrostatic interaction between two atoms. | [14] |
| CoulombEigFeat | Same as CulombFeat but it also calculates the eigenvalues of Coulomb matrices for molecules. | [14] |
| SmileImageFeat | The SmilesImageFeat featurizer transforms a SMILES string into an image. The default image size is 80 x 80, supporting two modes: std, a grayscale representation using atomic numbers for atom positions and a constant value for bonds, and engd, a 4-channel specification incorporating atom properties like hybridization, valency, charges, and bond type for enhanced visualization. Atom coordinates are computed, and lines between atoms indicate bonds, with | [15] |

| | | |
|---|---|---|
| | channels reflecting specified property values. | |
| SmilesSeqFeat | The SmilesSeqFeat featurizer converts a SMILES string into a sequence. SMILES below a specified maximum length are padded, while longer are excluded. The resulting sequence of character indices, obtained through a character-to-index mapping, can serve as input for a predictive model. | [15] |
| DMPNNFeat | This serves as a featurizer for the implementation of Directed Message Passing Neural Network (D-MPNN). The default node features include atomic number, degree, formal charge, chirality, number of hydrogens, hybridization, aromaticity, and mass, resulting in a feature length of 133. Edge features encompass bond type, same-ring membership, conjugation, and stereo configuration, with a feature length of 14. | [12, 16] |
| MatFeat | This functions as a featurizer for the Molecule Attention Transformer, producing a numpy array containing molecular graph descriptions, including node features, adjacency matrix, and distance matrix. | [17] |

Table S5 - Scikit-Learn scalers available in DeepMol

| Scaler | Description | References |
|---|---|---|
| StandardScaler | Standardize features by removing the mean and scaling to unit variance. | [18] |
| MinMaxScaler | Transform features by scaling each one of them to a given range. | [19] |
| RobustScaler | Scales features by subtracting the median and adjusting data based on the quantile range. This scaler is robust to outliers. | [20] |

| | | |
|---|---|---|
| PolynomialFeatures | Creates polynomial and interaction features by generating a new feature matrix that includes all polynomial combinations of the original features up to a specified degree. | [21] |
| Normalizer | Normalizes samples individually to unit norm. | [22] |
| Binarizer | Binarizes data (0 or 1) according to a threshold. | [23] |
| KernelCenterer | Centrally aligns kernel matrices by subtracting the mean along each feature dimension, ensuring that the kernel's diagonal elements are centered around zero. | [24] |
| QuantileTransformer | Adjusts features to follow a uniform or normal distribution, spreading out frequent values and mitigating the impact of outliers. | [25] |
| PowerTransformer | Applies a featurewise power transform to make the data more Gaussian-like. | [26] |

Table S6 - Scikit-Learn feature selection methods available in DeepMol

| Feature Selection Method | Description | References |
|---|---|---|
| LowVarianceFS | Removes all features with a variance lower than a threshold. | [27] |
| KbestFS | Selects the top k features based on their statistical significance with the target variable. It scores and retains the features with the highest correlation or dependency with the target. | [28] |
| PercentilFS | The PercentilFS method is a form of univariate feature selection that | [29] |

| | | |
|---|---|---|
| | involves choosing features through univariate statistical tests. It removes all features except for a user-defined highest-scoring percentage. | |
| RFECVFS | Selects features based on recursive feature elimination with cross-validation. It systematically eliminates less relevant features based on an estimator's performance through cross-validation to determine the optimal subset of features | [30] |
| SelectFromModelFS | It selects important features based on the coefficients or importance scores derived from a trained estimator. It allows for automatic feature selection by considering features that meet a specified threshold of importance. | [31] |

Table S7 - Data splitters in DeepMol - all of them can perform train-test, train-validation-test and k-fold splitting.

| Data Splitter | Description | References |
|---|---|---|
| RandomSplitter | Randomly splits the data. | - |
| SingletaskStratifiedSplitter | Splits single-task data on a stratified fashion. | |
| SimilaritySplitter | Splits molecules based on fingerprint similarity using Tanimoto similarity over Morgan fingerprints. It can create both homogeneous and heterogeneous splits. In homogeneous splits, molecules are evenly distributed across sets, whereas in heterogeneous splits, the goal is to minimize the similarity between molecules in one set and those in the other sets. | - |
| ScaffoldSplitter | Splits molecules according to the Bemis-Murcko scaffold representation, which represents rings, linkers, frameworks (combinations of linkers and | [32] |

| | | |
|---|---|---|
| | rings), and atomic properties like atom type, hybridization, and bond order within a molecular dataset. It can create both homogeneous and heterogeneous splits. In homogeneous splits, molecules are evenly distributed across sets, whereas in heterogeneous splits, the goal is to minimize the number of shared scaffolds between splits. | |
| ButinaSplitter | Splits the molecules based on the butina clustering of a bulk tanimoto fingerprint matrix. It can create both homogeneous and heterogeneous splits. In homogeneous splits, molecules are evenly distributed across sets, whereas in heterogeneous splits, molecules from the same clusters are kept in the same split. | [33] |
| MultiTaskStratifiedSplitter | Splits multi-task data on a stratified fashion. | - |

Table S8 - Imbalanced learning techniques provided by DeepMol.

| Unbalanced Learn Method | Description | References |
|---|---|---|
| Over-sampling techniques | | |
| RandomOverSampler | Randomly samples, with replacement, molecules in the under-represented class(es). | - |
| SMOTE | Synthetic Minority Oversampling Technique, is a statistical method aimed at balancing imbalanced datasets by creating new instances for the minority class(es). It generates new examples by combining features from existing minority cases and their nearest neighbors in the feature space. | [34] |

| Under-sampling techniques | | |
|---|---|---|
| RandomUnderSampler | Randomly under-samples, without replacement, molecules in the over-represented class(es). | - |
| ClusterCentroids | Under-samples data by generating centroids through clustering methods (KMeans algorithm by default). This algorithm keeps N majority samples using the coordinates of the N cluster centroids as the new majority samples. | - |
| Combination of over and under-sampling techniques | | |
| SMOTEENN | Technique that combines over- and under-sampling using SMOTE and Edited Nearest Neighbours (ENN) respectively. ENN removes examples for which the majority class label conflicts with the labels of the majority of its three nearest neighbors. | [35] |
| SMOTETomek | Technique that combines over- and under-sampling using SMOTE and Tomek links respectively. Tomek links identify pairs of instances, one from the majority class and one from the minority class, that are close to each other but have different class labels. The instances from the majority class are then removed. | [35] |

Table S9 - Scikit-learn, Deepchem and Keras models provided by DeepMol by default.

| Model | Description | References |
|---|---|---|
| Scikit-learn | | [36] |

| | | |
|---|---|---|
| 82 models as in scikit-learn (v1.2.0) | DeepMol includes all machine learning models available through Scikit-learn for regression, classification and multioutput. | - |
| DeepChem | | [37] |
| GatModel | A Graph Attention Network (GAT)-based model for predicting graph properties. It involves updating node representations using a GAT variant. The model computes the graph representation by combining weighted sums and max pooling of node representations, followed by concatenation, and utilizes a Multi-Layer Perceptron (MLP) for the final prediction. Works both for classification and regression. | [38] |
| GCNModel | Graph Convolution Networks (GCN)-based model for graph property prediction. It involves updating node representations using GCN. It then computes each graph's representation through a weighted sum and max pooling of node representations, followed by concatenation of the outputs. The final prediction is done using a Multilayer Perceptron (MLP). Works both for classification and regression. | [39] |
| AttentiveFPModel | Graph Property Prediction Model that involves combining node and edge features to initialize node representations through a round of message passing. Subsequently, node representations are updated with multiple rounds of message passing, and for each graph, its representation is computed by combining the representations of all nodes using a gated recurrent unit (GRU), followed by making the final prediction using a linear layer. Works both for classification and regression. | [40] |

| | | |
|---|---|---|
| PagtnModel | Graph Property Prediction model that employs a modified Graph Attention Network (GAT) to update node representations in graphs, utilizing a linear additive attention mechanism based on concatenating node and edge features. Multiple rounds of message passing are applied with residual connections between each layer, and the final molecular representation is obtained by aggregating node representations. The final prediction is obtained through a linear layer. Works both for classification and regression. | [41] |
| MPNNModel | Message Passing Neural Networks (MPNN) that considers graph convolutional operations as a specific form of a broader message passing scheme, wherein nodes exchange "messages", leading to updates in their internal states. The ordering of structures in this model adheres to the principles outlined in [43]. Works both for classification and regression. | [42, 43] |
| MEGNetModel | MatErials Graph Network uses multiple layers of Graph Networks known as MEGNetBlocks for predicting properties in molecules and crystals. It integrates node and edge properties through a Set2Set layer, combining this information with global features to perform property prediction tasks for materials or molecules. Works both for classification and regression. | [44] |
| DMPNNModel | Directed Message Passing Neural Network (D-MPNN), consisting of two phases: message-passing and read-out. In the message-passing phase, the objective is to generate hidden states for all atoms in the molecule using encoders, followed by the read-out phase where features are input into a feed-forward neural network to obtain task-based predictions. Works both for | [45] |

| | | |
|---|---|---|
| | classification and regression. | |
| CNN | 1, 2, or 3 dimensional Convolution Neural Network (CNN) comprising of a variable number of convolutional layers, followed by a global pooling layer (max pool or average pool), and a final fully connected layer for output computation. Works both for classification and regression. | - |
| MultitaskClassifier | A fully connected multitask classification network that offers extensive customization of the model, including options for adjusting the number and widths of layers, activation functions, regularization methods, and more. | - |
| MultitaskIRVClassifier | The IRV is a low-parameter neural network that enhances a k-nearest neighbor classifier by nonlinearly combining the impacts of neighboring chemicals in the training set. This model is used for multitask classification. | [46] |
| ProgressiveMultitaskClassifier | Progressive networks facilitate multitask learning by assigning a new set of weights to each task, preventing exponential forgetting and ensuring that prior tasks are not disregarded. This approach prevents the issue of exponential forgetting, ensuring the retention of knowledge from previous tasks in multitask learning. Used for multitask classification. | [47] |
| ProgressiveMultitaskRegressor | The same as ProgressiveMultitaskClassifier but for multitask regression. | [47] |
| RobustMultitaskClassifier | This model's fundamental concept involves incorporating bypass layers that directly connect features to the task output, offering | [48] |

| | | |
|---|---|---|
| | potential flexibility to navigate multitasking challenges affected by destructive interference. The inclusion of these bypass layers aims to facilitate adaptability in overcoming obstacles during multitasking. Used for multitask classification. | |
| RobustMultitaskRegressor | The same as RobustMultitaskClassifier but for multitask regression. | [48] |
| ScScoreModel | The SCScore model is a neural network that predicts the synthetic complexity score (SCScore) for molecules and establishes a correlation with the number of reaction steps needed to synthesize the target molecule. The model was adapted for classification tasks. | [49] |
| ChemCeption | The ChemCeption model applies convolutional neural networks (CNNs) to predict molecular properties by utilizing an image-based representation of the molecule. In this representation, various atomic and bond properties are encoded as pixels. Works both for classification and regression. | [50] |
| DAGModel | Directed Acyclic Graph models are used for predicting molecular properties by representing molecules as a series of directed graphs. In this approach, each atom in the molecule is transformed into a directed acyclic graph with edges pointing “inwards” to it. Works both for classification and regression. | [51] |
| GraphConvModel | Graph Convolutional Models, based on Duvenaud's convolutions. The model starts with per-atom descriptors for each molecule, undergoing a series of convolutional layers that combine and recombine these descriptors. | [10] |

| | | |
|---|---|---|
| Smiles2Vec | The Smiles2Vec model is implemented to generate neural representations of SMILES strings for subsequent tasks involving molecular properties. Using an Embedding layer, the model transforms input SMILES strings into vector representations, potentially incorporating a 1D convolutional layer and RNN cells to capture temporal dependencies and molecular structure information, facilitating molecular property prediction. Works both for classification and regression. | [52] |
| TextCNNModel | The model uses multiple 1D convolutional filters on padded strings, followed by max-over-time pooling, extracting individual features. After concatenating these features, the model undergoes transformations through hidden layers to make predictions. Works both for classification and regression. | [53] |
| WeaveModel | WeaveModel style convolutions differ from GraphConvModel style convolutions primarily in explicitly modeling bond features. This explicit modeling requires constructing an NxN matrix for bond interactions, potentially leading to scaling issues but offering the potential for more precise representation of subtle bond effects. Only for regression tasks. | [12] |
| DTNNModel | Deep Tensor Neural Network (DTNN), is a deep learning architecture designed specifically for understanding quantum-mechanical properties of molecular systems. DTNNs are particularly effective in capturing complex relationships within molecular structures, enabling insights into properties such as stability, atomic energies, and electronic structure. Only for regression tasks. | [54] |

| | | |
|---|---|---|
| MATModel | Model based on the Molecular Attention Transformer. It works by decomposing each molecule into its Node Features matrix, adjacency matrix, and distance matrix. Subsequently, a mask tensor is computed for the batch, serving as input for the MATEmbedding, MATEncoder, and MATGenerator layers. Works for regression tasks. | [55] |
| MultitaskRegressor | The same as MultitaskClassifier but for regression. | - |
| Tensorflow/Keras | | |
| FCNN | Fully connected neural network (FCNN) that allows flexible specification of the model architecture and training parameters. It supports multiple tasks, customizable hidden layers with activations, regularizers, and dropouts, along with options for batch normalization. It constructs a model with input, shared hidden layers, and task-specific output layers, compiling it with specified optimizer, loss functions, and evaluation metrics. Suitable for both classification and regression. | - |
| 1D CNN | 1D convolutional neural network (1D CNN) model suitable for classification and regression. It allows flexible specification of convolutional layers with options for filters, kernel sizes, activations, dropouts, and batch normalizations. The model architecture includes Gaussian noise injection, convolutional layers, dense layers, and task-specific output layers, compiled with specified optimizer, loss functions, and evaluation metrics. | - |
| TabularTransformer | Tabular transformer model based on the transformer architecture, including embedding layers, multi-head attention | - |

| | | |
|---|---|---|
| | layers, layer normalization, and dense layers. The model allows flexible specification of parameters such as attention layers, dropout rates, and activation functions for last layers, compiled with specified optimizer, loss functions, and evaluation metrics. Suitable for both classification and regression. | |
| RNN | This Recurrent Neural Network (RNN) model that supports flexible specification of parameters including the number of LSTM and GRU layers, units in each layer, dropout rates, and activation functions for both dense and last layers. The model is compiled with specified optimizer, loss functions, and evaluation metrics. Suitable for both classification and regression. | - |
| BidirectionalRNN | Bidirectional RNN model similar to the RNN model. However, it incorporates bidirectional LSTM and GRU layers, allowing the model to learn from both past and future information in the input sequences. It supports flexible specification of parameters such as the number of LSTM and GRU layers, units in each layer, dropout rates, and activation functions for both dense and last layers. The model is compiled with specified optimizer, loss functions, and evaluation metrics. Suitable for both classification and regression. | - |

Table S10 - Feature explainability using Shapley values (SHAP [56]) in DeepMol.

| Explainer | Description |
|---|---|
| Permutation | Approximates SHAP values by systematically permuting input features. |

| | |
|---|---|
| Exact | Computes SHAP values by using optimized exact enumeration techniques. It is particularly suited for models with less than 15 features. |
| Additive | Computes SHAP values for generalized additive models. |
| Tree | Computes SHAP values for ensemble tree models. |
| GPU Tree | GPU accelerated version of TreeExplainer. |
| Partition | Recursively computes SHAP values through a hierarchy of features, accounting for correlations among features by grouping them together |
| Linear | Computes SHAP values for linear models. |
| Sampling | Computes SHAP values under the assumption of feature independence. |
| Deep | Approximates SHAP values for deep learning models. |
| Kernel | Computes SHAP values by using a weighted linear regression to compute the importance of each feature. |
| Random | Returns random (normally distributed) feature attributions. Only for benchmark comparisons. |

### Voting pipelines and ensembles

Table S11 - Ensembles and voting pipelines

| Dataset | Ensemble type | Models/Pipelines |
|---|---|---|
| Bioav | Voting Pipeline | • 5 ChEMBL Standardizer + DMPNN |
| Lipo | Stacking Ensemble | • Linear Regression<br>• SVM<br>• Random Forest<br>• Gradient Boosting<br>• Final estimator: MLP |
| BBB | Voting Pipeline | • 3 Custom Standardizer + DMPNN<br>• ChEMBL Standardizer + DMPNN<br>• ChEMBL Standardizer + GCN |
| PPBR | Stacking Ensemble | • Linear Regression<br>• SVM<br>• Random Forest<br>• Gradient Boosting<br>• Final estimator: MLP |
| CYP2C9 Inhibition | Voting Pipeline | • Basic Standardizer + MorganFingerprint + SVC<br>• Custom Standardizer + AtomPairFingerprint + LowVarianceFS + GradientBoosting<br>• BasicStandardizer + MorganFingerprint + GradientBoosting<br>• BasicStandardizer + MorganFingerprint + RidgeClassifierCV<br>• AtomPairFingerprint + LowVarianceFS + GradientBoosting |
| CYP2C9 Substrate | Bagging Ensemble | 150 Logistic Regression |
| CYP2D6 Inhibition | Stacking Ensemble | • LogisticRegression<br>• SVC<br>• RandomForest<br>• GradientBoosting<br>• Final estimator: MLP |
| CYP3A4 Inhibition | Voting Pipeline | • ChEMBL Standardizer + ConvMolFeat + GraphConvModel |

| | | |
|---|---|---|
| | | • 4 Basic Standardizer + ConvMolFeat + GraphConvModel |
| CYP3A4 Substrate | Bagging Ensemble | 450 SVMs |
| CL-Hepa | Voting Pipeline | • 3 ChEMBL Standardizer + DMPNN<br>• GCN<br>• ChEMBL Standardizer + GCN |
| CL-Micro | Voting Pipeline | 5 ChEMBL Standardizer + TextCNN |
| Ames | Voting Pipeline | 5 ChEMBL Standardizer + GCN |
| LD50 | Voting Regressor | • Linear Regression<br>• SVM<br>• Random Forest<br>• Gradient Boosting<br>• MLP |
| DILI | Voting Pipeline | • Basic Standardizer + Layered and Morgan FPs + 1D CNN<br>• 4 Custom Standardizers + MACCS keys + Select from model + Gaussian Process classifier |

### Comparison with similar tools

In recent years, several chemoinformatics toolkits, including DeepChem, OpenChem, AMPL and ZairaChem have been developed. These open-source projects, written in Python, are utilized for machine learning in constructing pipelines for QSAR/QSPR systems. Regarding dataset loading, DeepMol and DeepChem support CSV and SDF formats, while OpenChem, AMPL and ZairaChem only handle CSV files, which limits their capability to import pre-computed 3D structures.

DeepMol offers various methods for molecule standardization, whereas OpenChem allows sanitization, padding sequences, and canonizing SMILES, and AMPL enables striping salt groups, neutralizing, kekulizing molecules, and replacing rare isotopes with the most common ones for each element. Only DeepChem supports molecule sanitization during dataset loading. ZairaChem applies the MELLODDY-Tuner protocol, which includes disconnecting metal atoms from the rest of the molecule, reionizing the molecule, adjusting charges and protonation states, assigning stereochemistry to the molecule, ensuring consistent representation of chiral centres, and additional standardization procedures. All tools support data splitting, except OpenChem, and only DeepMol can perform a stratified split specifically designed for molecular data and multitask classification.

Additionally, all the toolkits can generate numerical features from molecules. While these toolkits primarily focus on deep learning models for model construction, they all provide shallow learning models except for OpenChem. Hyperparameter and architecture optimization are possible in all the toolkits except for OpenChem. Furthermore, although all methods offer pipelines, DeepMol and ZairaChem stand out by providing a comprehensive and efficient approach to building, testing, optimizing, and deploying pipelines. DeepMol excels compared to other tools in providing feature importance analysis and effectively connecting features to molecular structures. It also offers methods to handle unbalanced datasets and perform integrated and automatic feature selection, distinguishing it from other assessed tools.

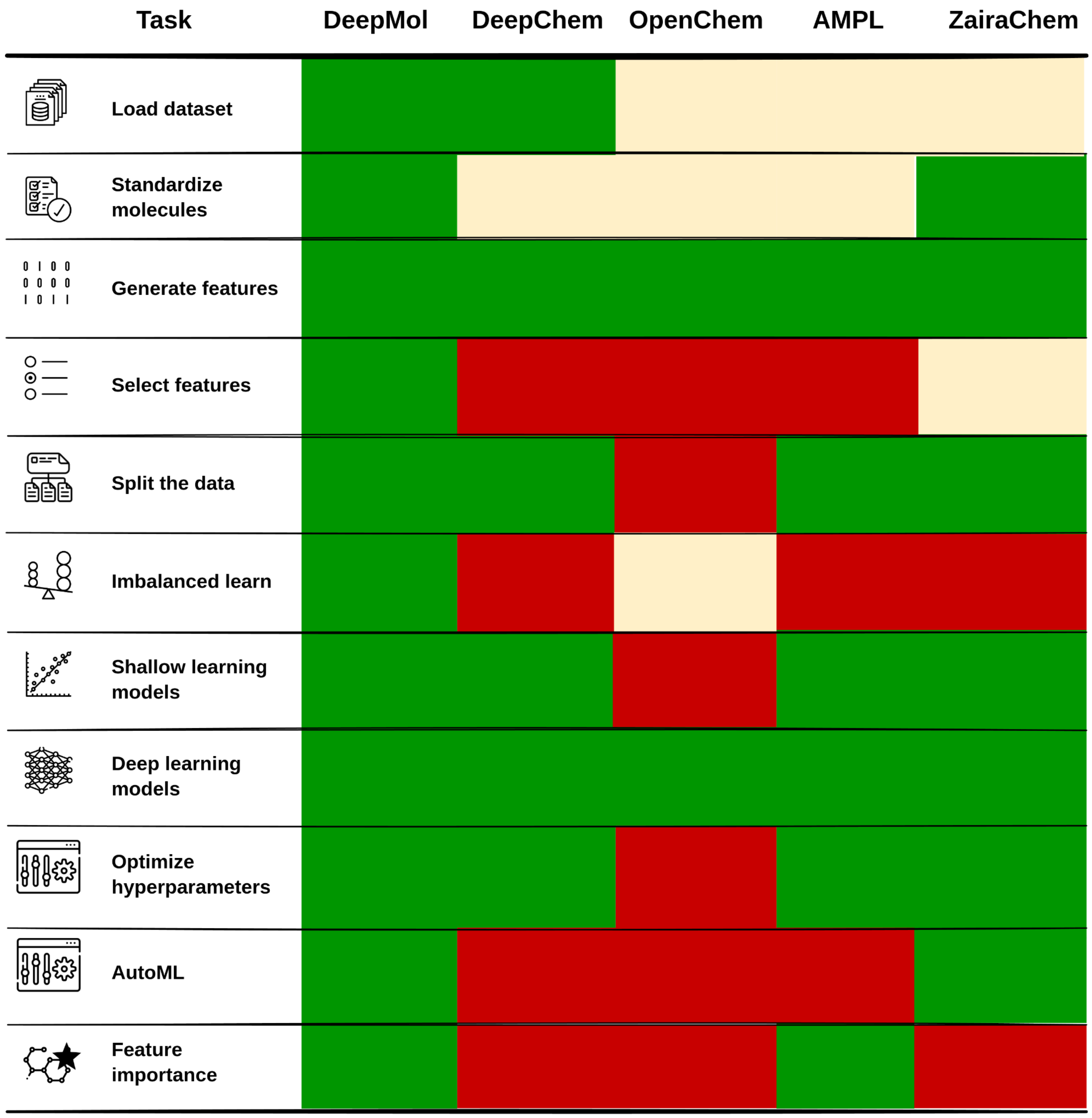


Figure S1 - Heatmap showcasing the presence or absence of an integrated way of performing relevant tasks for constructing QSAR/QSPR systems. Green stands for presence, red for absence and yellow for incomplete integration.

References:

[1] Rogers, D., & Hahn, M. (2010). Extended-Connectivity Fingerprints. In Journal of Chemical Information and Modeling (Vol. 50, Issue 5, pp. 742–754). American Chemical Society (ACS). https://doi.org/10.1021/ci100050t

[2] Durant, J. L., Leland, B. A., Henry, D. R., & Nourse, J. G. (2002). Reoptimization of MDL Keys for Use in Drug Discovery. In Journal of Chemical Information and Computer Sciences (Vol. 42, Issue 6, pp. 1273–1280). American Chemical Society (ACS). https://doi.org/10.1021/ci010132r
[3] https://www.daylight.com/dayhtml/doc/theory/theory.finger.html
[4] Carhart, R. E., Smith, D. H., & Venkataraghavan, R. (1985). Atom pairs as molecular features in structure-activity studies: definition and applications. In Journal of Chemical Information and Computer Sciences (Vol. 25, Issue 2, pp. 64–73). American Chemical Society (ACS). https://doi.org/10.1021/ci00046a002
[5] Todeschini, R., & Consonni, V. (2003). Descriptors from Molecular Geometry. In Handbook of Chemoinformatics (pp. 1004–1033). Wiley. https://doi.org/10.1002/9783527618279.ch37
[6] Firth, N. C., Brown, N., & Blagg, J. (2012). Plane of Best Fit: A Novel Method to Characterize the Three-Dimensionality of Molecules. In Journal of Chemical Information and Modeling (Vol. 52, Issue 10, pp. 2516–2525). American Chemical Society (ACS). https://doi.org/10.1021/ci300293f
[7] Arteca, G. A. (1996). Molecular Shape Descriptors. In Reviews in Computational Chemistry (pp. 191–253). Wiley. https://doi.org/10.1002/9780470125861.ch5
[8] Baumgärtner, A. (1993). Shapes of flexible vesicles at constant volume. In The Journal of Chemical Physics (Vol. 98, Issue 9, pp. 7496–7501). AIP Publishing. https://doi.org/10.1063/1.464689
[9] Sauer, W. H. B., & Schwarz, M. K. (2003). Molecular Shape Diversity of Combinatorial Libraries:  A Prerequisite for Broad Bioactivity. In Journal of Chemical Information and Computer Sciences (Vol. 43, Issue 3, pp. 987–1003). American Chemical Society (ACS). https://doi.org/10.1021/ci025599w
[10] Duvenaud, D. K., Maclaurin, D., Iparraguirre, J., Bombarell, R., Hirzel, T., Aspuru-Guzik, A. & Adams, R. P. (2015). Convolutional Networks on Graphs for Learning Molecular Fingerprints. In C. Cortes, N. D. Lawrence, D. D. Lee, M. Sugiyama & R. Garnett (ed.), Advances in Neural Information Processing Systems 28 (pp. 2224--2232) . Curran Associates, Inc. .
[11] Chen, B., Barzilay, R., & S Jaakkola, T. (2019). Path-Augmented Graph Transformer Network. American Chemical Society (ACS). https://doi.org/10.26434/chemrxiv.8214422.v1
[12] Kearnes, S., McCloskey, K., Berndl, M., Pande, V., & Riley, P. (2016). Molecular graph convolutions: moving beyond fingerprints. In Journal of Computer-Aided Molecular Design (Vol. 30, Issue 8, pp. 595–608). Springer Science and Business Media LLC. https://doi.org/10.1007/s10822-016-9938-8

[13] De Cao, N., & Kipf, T. (2018). MolGAN: An implicit generative model for small molecular graphs. arXiv. https://doi.org/10.48550/ARXIV.1805.11973
[14] Montavon, G., Hansen, K., Fazli, S., Rupp, M., Biegler, F., Ziehe, A., … Müller, K.-R. (2012). Learning Invariant Representations of Molecules for Atomization Energy Prediction. In F. Pereira, C. J. Burges, L. Bottou, & K. Q. Weinberger (Eds.), Advances in Neural Information Processing Systems (Vol. 25).
[15] Goh, G. B., Siegel, C., Vishnu, A., & Hodas, N. (2018). Using Rule-Based Labels for Weak Supervised Learning. In Proceedings of the 24th ACM SIGKDD International Conference on Knowledge Discovery & Data Mining. KDD '18: The 24th ACM SIGKDD International Conference on Knowledge Discovery and Data Mining. ACM. https://doi.org/10.1145/3219819.3219838
[16] Yang, K., Swanson, K., Jin, W., Coley, C., Eiden, P., Gao, H., Guzman-Perez, A., Hopper, T., Kelley, B., Mathea, M., Palmer, A., Settels, V., Jaakkola, T., Jensen, K., & Barzilay, R. (2019). Analyzing Learned Molecular Representations for Property Prediction. In Journal of Chemical Information and Modeling (Vol. 59, Issue 8, pp. 3370–3388). American Chemical Society (ACS). https://doi.org/10.1021/acs.jcim.9b00237
[17] Maziarka, Ł., Danel, T., Mucha, S., Rataj, K., Tabor, J., & Jastrzębski, S. (2020). Molecule Attention Transformer. arXiv. https://doi.org/10.48550/ARXIV.2002.08264
[18] Sklearn.preprocessing.StandardScaler. scikit. (n.d.). https://scikit-learn.org/stable/modules/generated/sklearn.preprocessing.StandardScaler.html#sklearn.preprocessing.StandardScaler
[19] Sklearn.preprocessing.MinMaxScaler. scikit. (n.d.-a). https://scikit-learn.org/stable/modules/generated/sklearn.preprocessing.MinMaxScaler.html#sklearn.preprocessing.MinMaxScaler
[20] Sklearn.preprocessing.RobustScaler. scikit. (n.d.-b). https://scikit-learn.org/stable/modules/generated/sklearn.preprocessing.RobustScaler.html#sklearn.preprocessing.RobustScaler
[21] Sklearn.preprocessing.PolynomialFeatures. scikit. (n.d.-b). https://scikit-learn.org/stable/modules/generated/sklearn.preprocessing.PolynomialFeatures.html#sklearn.preprocessing.PolynomialFeatures
[22] Sklearn.preprocessing.Normalizer. scikit. (n.d.-b). https://scikit-learn.org/stable/modules/generated/sklearn.preprocessing.Normalizer.html#sklearn.preprocessing.Normalizer
[23] Sklearn.preprocessing.Binarizer. scikit. (n.d.-a). https://scikit-learn.org/stable/modules/generated/sklearn.preprocessing.Binarizer.html#sklearn.preprocessing.Binarizer

[24] Sklearn.preprocessing.KernelCenterer. scikit. (n.d.-b). https://scikit-learn.org/stable/modules/generated/sklearn.preprocessing.KernelCenterer.html#sklearn.preprocessing.KernelCenterer
[25] Sklearn.preprocessing.QuantileTransformer. scikit. (n.d.-f). https://scikit-learn.org/stable/modules/generated/sklearn.preprocessing.QuantileTransformer.html#sklearn.preprocessing.QuantileTransformer
[26] Sklearn.preprocessing.PowerTransformer. scikit. (n.d.-f). https://scikit-learn.org/stable/modules/generated/sklearn.preprocessing.PowerTransformer.html#sklearn.preprocessing.PowerTransformer
[27] Sklearn.feature_selection.Variancethreshold. scikit. (n.d.-a). https://scikit-learn.org/stable/modules/generated/sklearn.feature_selection.VarianceThreshold.html#sklearn.feature_selection.VarianceThreshold
[28] Sklearn.feature_selection.Selectkbest. scikit. (n.d.-a). https://scikit-learn.org/stable/modules/generated/sklearn.feature_selection.SelectKBest.html#sklearn.feature_selection.SelectKBest
[29] Sklearn.feature_selection.Selectpercentile. scikit. (n.d.-b). https://scikit-learn.org/stable/modules/generated/sklearn.feature_selection.SelectPercentile.html#sklearn.feature_selection.SelectPercentile
[30] Sklearn.feature_selection.RFECV. scikit. (n.d.-a). https://scikit-learn.org/stable/modules/generated/sklearn.feature_selection.RFECV.html#sklearn.feature_selection.RFECV
[31] Sklearn.feature_selection.Selectfrommodel. scikit. (n.d.-b). https://scikit-learn.org/stable/modules/generated/sklearn.feature_selection.SelectFromModel.html#sklearn.feature_selection.SelectFromModel
[32] Bemis, G. W., & Murcko, M. A. (1996). The Properties of Known Drugs. 1. Molecular Frameworks. In Journal of Medicinal Chemistry (Vol. 39, Issue 15, pp. 2887–2893). American Chemical Society (ACS). https://doi.org/10.1021/jm9602928
[33] Butina, D. (1999). Unsupervised Data Base Clustering Based on Daylight's Fingerprint and Tanimoto Similarity: A Fast and Automated Way To Cluster Small and Large Data Sets. In Journal of Chemical Information and Computer Sciences (Vol. 39, Issue 4, pp. 747–750). American Chemical Society (ACS). https://doi.org/10.1021/ci9803381
[34] Chawla, N. V., Bowyer, K. W., Hall, L. O., & Kegelmeyer, W. P. (2002). SMOTE: Synthetic Minority Over-sampling Technique. In Journal of Artificial Intelligence Research (Vol. 16, pp. 321–357). AI Access Foundation. https://doi.org/10.1613/jair.953
[35] Batista, G. E. A. P. A., Prati, R. C., & Monard, M. C. (2004). A study of the behavior of several methods for balancing machine learning training data. In ACM SIGKDD Explorations Newsletter (Vol. 6, Issue 1, pp. 20–29).

Association for Computing Machinery (ACM).
https://doi.org/10.1145/1007730.1007735
[36] Fabian Pedregosa, Gael Varoquaux, Alexandre Gramfort, Vincent Michel, Bertrand Thirion, Olivier Grisel, Mathieu Blondel, Peter Prettenhofer, Ron Weiss, Vincent Dubourg, Jake Vanderplas, Alexandre Passos, David Cournapeau, Matthieu Brucher, Matthieu Perrot, & Edouard Duchesnay (2011). Scikit-learn: Machine Learning in Python. Journal of Machine Learning Research, 12(85), 2825-2830.
[37] Bharath Ramsundar, Peter Eastman, Patrick Walters, Vijay Pande, Karl Leswing, & Zhenqin Wu (2019). Deep Learning for the Life Sciences. O'Reilly Media.
[38] Veličković, P., Cucurull, G., Casanova, A., Romero, A., Liò, P., & Bengio, Y. (2017). Graph Attention Networks (Version 3). arXiv.
https://doi.org/10.48550/ARXIV.1710.10903
[39] Kipf, T. N., & Welling, M. (2016). Semi-Supervised Classification with Graph Convolutional Networks (Version 4). arXiv.
https://doi.org/10.48550/ARXIV.1609.02907
[40] Xiong, Z., Wang, D., Liu, X., Zhong, F., Wan, X., Li, X., Li, Z., Luo, X., Chen, K., Jiang, H., & Zheng, M. (2019). Pushing the Boundaries of Molecular Representation for Drug Discovery with the Graph Attention Mechanism. In Journal of Medicinal Chemistry (Vol. 63, Issue 16, pp. 8749–8760). American Chemical Society (ACS). https://doi.org/10.1021/acs.jmedchem.9b00959
[41] Chen, B., Barzilay, R., & Jaakkola, T. (2019). Path-Augmented Graph Transformer Network (Version 1). arXiv.
https://doi.org/10.48550/ARXIV.1905.12712
[42] Gilmer, J., Schoenholz, S. S., Riley, P. F., Vinyals, O., & Dahl, G. E. (2017). Neural Message Passing for Quantum Chemistry (Version 2). arXiv.
https://doi.org/10.48550/ARXIV.1704.01212
[43] Vinyals, O., Bengio, S., & Kudlur, M. (2015). Order Matters: Sequence to sequence for sets (Version 4). arXiv.
https://doi.org/10.48550/ARXIV.1511.06391
[44] Chen, C., Ye, W., Zuo, Y., Zheng, C., & Ong, S. P. (2018). Graph Networks as a Universal Machine Learning Framework for Molecules and Crystals. arXiv. https://doi.org/10.48550/ARXIV.1812.05055
[45] Yang, K., Swanson, K., Jin, W., Coley, C., Eiden, P., Gao, H., Guzman-Perez, A., Hopper, T., Kelley, B., Mathea, M., Palmer, A., Settels, V., Jaakkola, T., Jensen, K., & Barzilay, R. (2019). Analyzing Learned Molecular Representations for Property Prediction. In Journal of Chemical Information and Modeling (Vol. 59, Issue 8, pp. 3370–3388). American Chemical Society (ACS). https://doi.org/10.1021/acs.jcim.9b00237

[46] Swamidass, S. J., Azencott, C.-A., Lin, T.-W., Gramajo, H., Tsai, S.-C., & Baldi, P. (2009). Influence Relevance Voting: An Accurate And Interpretable Virtual High Throughput Screening Method. In Journal of Chemical Information and Modeling (Vol. 49, Issue 4, pp. 756–766). American Chemical Society (ACS). https://doi.org/10.1021/ci8004379
[47] Rusu, A. A., Rabinowitz, N. C., Desjardins, G., Soyer, H., Kirkpatrick, J., Kavukcuoglu, K., Pascanu, R., & Hadsell, R. (2016). Progressive Neural Networks (Version 4). arXiv. https://doi.org/10.48550/ARXIV.1606.04671
[48] Ramsundar, B., Liu, B., Wu, Z., Verras, A., Tudor, M., Sheridan, R. P., & Pande, V. (2017). Is Multitask Deep Learning Practical for Pharma? In Journal of Chemical Information and Modeling (Vol. 57, Issue 8, pp. 2068–2076). American Chemical Society (ACS). https://doi.org/10.1021/acs.jcim.7b00146
[49] Coley, C. W., Rogers, L., Green, W. H., & Jensen, K. F. (2018). SCScore: Synthetic Complexity Learned from a Reaction Corpus. In Journal of Chemical Information and Modeling (Vol. 58, Issue 2, pp. 252–261). American Chemical Society (ACS). https://doi.org/10.1021/acs.jcim.7b00622
[50] Goh, G. B., Siegel, C., Vishnu, A., Hodas, N. O., & Baker, N. (2017). Chemception: A Deep Neural Network with Minimal Chemistry Knowledge Matches the Performance of Expert-developed QSAR/QSPR Models (Version 1). arXiv. https://doi.org/10.48550/ARXIV.1706.06689
[51] Lusci, A., Pollastri, G., & Baldi, P. (2013). Deep Architectures and Deep Learning in Chemoinformatics: The Prediction of Aqueous Solubility for Drug-Like Molecules. In Journal of Chemical Information and Modeling (Vol. 53, Issue 7, pp. 1563–1575). American Chemical Society (ACS). https://doi.org/10.1021/ci400187y
[52] Goh, G. B., Hodas, N. O., Siegel, C., & Vishnu, A. (2017). SMILES2Vec: An Interpretable General-Purpose Deep Neural Network for Predicting Chemical Properties (Version 2). arXiv. https://doi.org/10.48550/ARXIV.1712.02034
[53] Guimaraes, G. L., Sanchez-Lengeling, B., Outeiral, C., Farias, P. L. C., & Aspuru-Guzik, A. (2017). Objective-Reinforced Generative Adversarial Networks (ORGAN) for Sequence Generation Models (Version 3). arXiv. https://doi.org/10.48550/ARXIV.1705.10843
[54] Schütt, K. T., Arbabzadah, F., Chmiela, S., Müller, K. R., & Tkatchenko, A. (2017). Quantum-chemical insights from deep tensor neural networks. In Nature Communications (Vol. 8, Issue 1). Springer Science and Business Media LLC. https://doi.org/10.1038/ncomms13890
[55] Maziarka, Ł., Danel, T., Mucha, S., Rataj, K., Tabor, J., & Jastrzębski, S. (2020). Molecule Attention Transformer. arXiv. https://doi.org/10.48550/ARXIV.2002.08264

[56] Lundberg, S., & Lee, S.-I. (2017). A Unified Approach to Interpreting Model Predictions (Version 2). arXiv. https://doi.org/10.48550/ARXIV.1705.07874
